# Pediatric nasal epithelial cells are less permissive to SARS-CoV-2 replication compared to adult cells

**DOI:** 10.1101/2021.03.08.434300

**Authors:** Yanshan Zhu, Keng Yih Chew, Melanie Wu, Anjana C. Karawita, Georgina McCallum, Lauren E Steele, Ayaho Yamamoto, Larisa L. Labzin, Tejasri Yarlagadda, Alexander A. Khromykh, Xiaohui Wang, Julian Sng, Claudia J. Stocks, Yao Xia, Tobias R. Kollmann, David Martino, Merja Joensuu, Frédéric A. Meunier, Giuseppe Balistreri, Helle Bielefeldt-Ohmann, Asha C. Bowen, Anthony Kicic, Peter D. Sly, Kirsten M. Spann, Kirsty R. Short

## Abstract

Children typically experience more mild symptoms of COVID-19 when compared to adults. There is a strong body of evidence that children are also less susceptible to SARS-CoV-2 infection with the ancestral viral isolate. However, the emergence of SARS-CoV-2 variants of concern (VOCs) has been associated with an increased number of pediatric infections. Whether this is the result of widespread adult vaccination or fundamental changes in the biology of SARS-CoV-2 remains to be determined. Here, we use primary nasal epithelial cells from children and adults, differentiated at an air-liquid interface to show that the ancestral SARS-CoV-2 replicates to significantly lower titers in the nasal epithelial cells of children compared to those of adults. This was associated with a heightened antiviral response to SARS-CoV-2 in the nasal epithelial cells of children. Importantly, the Delta variant also replicated to significantly lower titres in the nasal epithelial cells of children. This trend was markedly less pronounced in the case of Omicron. It is also striking to note that, at least in terms of viral RNA, Omicron replicated better in pediatric NECs compared to both Delta and the ancestral virus. Taken together, these data show that the nasal epithelium of children supports lower infection and replication of ancestral SARS-CoV-2, although this may be changing as the virus evolves.

## Introduction

Severe Acute Respiratory Syndrome-Coronavirus 2 (SARS-CoV-2), the causative agent of coronavirus disease-2019 (COVID-19), causes a broad range of clinical symptoms, ranging from asymptomatic infection to potentially fatal acute respiratory distress syndrome (ARDS). Children typically experience mild symptoms of COVID-19 when compared to adults [1]. There is also a significant body of evidence with the ancestral viral strain that children are less susceptible to SARS-CoV-2 infection and less likely to transmit the virus[2]. These findings have been echoed in multiple single site studies where, both within and outside of households, the infection rate of the ancestral SARS-CoV-2 amongst children <10 years old is significantly lower than that of adults [3]. Reduced SARS-CoV-2 infection and transmission is also observed in juvenile ferrets compared to their older counterparts [4].

The reasons for less frequent SARS-CoV-2 infection and symptoms in children infected with the ancestral virus strain remain unclear and may be influenced by a multitude of factors. There is evidence to suggest that nasal epithelial cells (NECs), the first site of infection, are fundamentally different in children compared to adults. Gene expression studies using the nasal epithelium of healthy individuals suggest that the transcript for the SARS-CoV-2 receptor, angiotensin converting enzyme-2 (*ACE2*), is expressed at lower levels in children than in adults [5]. However, this has yet to be validated on a protein level. Moreover, this does not appear to be the case in all patient cohorts [6, 7]. Following binding of the SARS-CoV-2 spike protein to ACE2, the host surface transmembrane serine protease 2 (TMPRSS2) is also involved in viral entry into the cell [8]. NECs from children express less *TMPRSS2* mRNA than those from adults, which may contribute to less frequent pediatric infections with SARS-CoV-2 [9]. However, this has also yet to be confirmed at protein level. In addition to differential receptor expression, pediatric and adult NECs may also mount fundamentally different innate immune response to SARS-CoV-2. Recent RNA sequencing of the whole epithelium from pediatric and adult proximal airways suggests that there is a higher expression of genes associated with inflammation and the anti-viral response in children compared to adults [10, 11]. Whilst increased inflammation and interferon production have previously been associated with elevated COVID-19 severity [12], it is important to note that such studies refer to the inflammatory response in the lower respiratory tract, where any immunopathology may lead to respiratory distress [13]. In contrast, inflammation in the upper respiratory tract plays an important role in controlling early viral replication. Specifically, elevated levels of type I IFN effectively inhibit the replication of SARS-CoV-2 across multiple studies [14-16]. Consistent with these data, nasopharyngeal swabs from SARS-CoV-2 infected children display elevated levels of interferons and inflammatory markers compared to those of SARS-CoV-2 infected adults [6].

During the course of the SARS-CoV-2 pandemic the ancestral virus has undergone significant mutations resulting in the emergence of variants of concern (VOCs) such as Delta and Omicron. These VOCs have multiple mutations in the spike protein, the N protein and various open reading frames (ORFs) of the virus, which has resulted in their increased transmissibility and potentially differing clinical outcome [17]. The growing dominance of VOCs has raised speculation that the epidemiology of SARS-CoV-2 infection has fundamentally changed, specifically in terms of the role that children play in spreading the virus. For example, epidemiological household transmission studies have found little evidence of differential susceptibility to Delta in children compared to adults [18, 19]. In addition, data from South Africa [20] and the US [21] found a rapid increase in pediatric COVID-19-related hospital admission associated with the Omicron wave. These data may suggest that VOCs have evolved such that they are now able to evade any protection that the innate immune response has previously afforded children in terms of infection with the ancestral virus [6, 11, 22, 23]. However, the extrapolation of epidemiology studies to fundamental immunology are complicated by the fact that the Delta and Omicron waves emerged at a time when a large percentage of the adult population were eligible for vaccination whilst vaccination from children <12 years old often lagged behind. Therefore, the role of pediatric innate immunity during VOC infection remains undefined.

Here, we use primary nasal epithelial cells (NECs), differentiated at an air-liquid interface, to investigate differential infection kinetics and antiviral responses to SARS-CoV-2 (ancestral and VOCs) infection in children and adults.

## Materials and methods

### Cell collection and ethics statement

Primary NECs were collected from healthy adult (aged 19 to 65 years old) donors by placing a sterile nasal mucosal curette (Arlington Scientific Inc., USA) in the mid-inferior portion of the inferior turbinate during July 2018 to May 2021 (during which time there was only sporadic community cases of COVID-19 in Queensland and Western Australia). Informed consent was obtained from all donors. Primary NECs were obtained from healthy pediatric donors (aged 2 to 11 years old) in the same manner while under general anesthetic prior to Ear-Nose and Throat (ENT)-related surgeries including tonsillectomies, adenoidectomies or for sleep apnoea. Children did not have any other unknown underlying condition at the time of recruitment and sampling. This study was approved by the University of Queensland’s Human Research Ethics Committee (2020001742), the Queensland Children’s Hospital and Health Service Human Research Ethics Committee (HREC/16/QRCH/215, HREC/10/QRCH/78), Queensland University of Technology Human Research Ethics Committee 17000000039), and the St John of God Subiaco Hospital Human Ethics Committee as part of the Western Australia Epithelial Research Program (WAERP) (Ethics #901). Primary NECs were established as previously described[24-26] and stored in freezing media (FBS with 10% DMSO) at passage 1 or 2. In total, we included 38 donors in this study(adult (N=15, 7 females, 8 male, 37.9±16.5 years old) and pediatric (N=23, 11 female, 12 male, 5.6±2.7 years old)).

### Cell culture

African green monkey kidney epithelial Vero cells were maintained in MEM (Invitrogen), containing 10% (v/v) heat-inactivated fetal bovine serum (Cytiva), 100 U/ml penicillin and streptomycin (Life Technologies Australia). Cell lines were obtained from American Type Culture Collection (ATCC; Virginia, USA). Primary NECs were expanded and passaged in Pneumacult EX Plus media (STEMCELL Technologies Inc, Canada). After initial expansion, NECs were seeded at a density of 4-5×10^5^ cells/transwell on 6.5 mm transwell polyester membranes with 0.4um pores (Corning Costar, USA) and cultured in EX Plus media (STEMCELL Technologies). Cells were monitored for confluence. When a confluent monolayer was achieved, cells were ‘air-lifted’ by removing the media from the apical chamber and replacing the basolateral media with Pneumacult air liquid interface (ALI) media (STEMCELL Technologies) [27]. Medium was replaced in the basal compartment three times a week, and the cells were maintained in ALI conditions for at least 3 weeks until ciliated cells and mucus were observed and cells obtained a transepithelial electrical resistance (TEER) measurement greater than 1000Ω. Fully differentiated cultures were used in downstream infection experiments using influenza virus or SARS-CoV-2.

### Viral stocks

SARS-CoV-2 isolate hCoV-19/Australia/QLD02/2020 (QLD02) (used as the original ancestral virus), hCoV-19/Australia/QLD1893C/2021 (QLD1893C) (GISAID Accession ID; EPI_ISL_2433928; Delta), and hCoV-19/Australia/NSW-RPAH-1933/2021 (GISAID Accession ID; EPI_ISL_6814922; Omicron) were kindly provided by Queensland Health Forensic & Scientific Services, Queensland Department of Health and the Kirby Institute. Virus was amplified in Vero cells expressing human TMPRSS2 and titrated by plaque assay [28]. All studies with SARS-CoV-2 were performed under physical containment 3 (PC3) conditions and were approved by the University of Queensland Biosafety Committee (IBC/374B/SCMB/2020).

### Viral infection

Differentiated adult and pediatric NECs were infected with mock (PBS), QLD02 (1.25 × 10^5^ PFU or 1.7 × 10^4^ PFU), QLD1517 (1.7 × 10^4^ PFU) or NSW1933 (1.7 × 10^4^ PFU). Specifically, 100uL of virus or PBS was placed on the epithelial surface in the apical compartment and incubated for 1 hour at 37ºC. Following incubation, excess virus was removed from the transwell and cells were incubated at 37ºC with 5% CO_2_. Every 24 hours the basolateral media was refreshed with 1mL of new ALI media. At pre-determined timepoints post-infection 100μL of PBS was added to the apical compartment and cells were incubated at 37ºC with 5% CO_2_ for 10 minutes. The apical supernatant was subsequently collected and stored at -80ºC. Cells were lysed with Buffer RLT (Qiagen, USA) containing 0.01% β-mercaptoethanol for RNA analysis. Alternatively, cells were lysed in 2% SDS/PBS lysis buffer (2% SDS/PBS buffer, 10% 10x PhosSTOP, 4% 25x protease inhibitor) for protein analysis or fixed overnight in 4% paraformaldehyde for histology.

### Histology

Fixed cells on a transwell membrane were routine processed and embedded in paraffin, sectioned at 5μm and subsequently stained with hematoxylin and eosin (H&E) or Periodic acid–Schiff (PAS). Sections were assessed for cellular morphology by a veterinary pathologist (H.B.O.) blinded to the experimental design.

### Immunofluorescence

Differentiated epithelial cells grown on a transwell membrane were fixed with 4% paraformaldehyde (Cat#15710, Electron Microscopy Sciences) in PBS for 45 minutes at room temperature, followed by a blocking with 0.5% BSA (Sigma) in PBS for 30 minutes and permeabilization with 0.02% of Triton X-100 (Sigma) in PBS for 15 min at room temperature. After washing twice with PBS/BSA and a second blocking step for 10 min at room temperature, samples were incubated with primary antibodies overnight at 4°C. Primary antibodies were diluted in 0.5% BSA in PBS blocking solution: 1:400 ZO-1 (Cat#40-2200, Thermo Fisher Scientific); 1:1000 MUC5AC (Cat#MA5-12178, ThermoFisher Scientific); 1:500 ACE2 (Cat#AF933, R&D Systems). After three washing steps with 0.5% BSA/PBS for 5 minutes each time, the samples were incubated in secondary antibody: 1:1000 Alexa Flour 555 donkey anti-goat (Cat#A21432, Invitrogen) for 2.5 hours at room temperature in dark, and after three washes in PBS and three washes with 0.5% BSA/PBS, the cells were incubated with a 1:1000 Alexa Fluor 488 goat anti-mouse (Cat#A32728, Invitrogen) for 2.5 hours at room temperature covered from light. The cells were simultaneously stained with 1:400 Alexa Fluor 647 Phalloidin (Cat#A22287 Invitrogen) and 1:1000 DAPI. After three washes in PBS, the transwell membranes with cells were cut with a scalpel, briefly dipped in milli-q water, and mounted on a class slide using ProLong Gold Antifade Mountant (Cat# P10144, ThermoFisher Scientific). Mounted samples were imaged on a spinning-disk confocal system (Marianas; 3I, Inc.) consisting of a Axio Observer Z1 (Carl Zeiss) equipped with a CSU-W1 spinning-disk head (Yokogawa Corporation of America), ORCA-Flash4.0 v2 sCMOS camera (Hamamatsu Photonics), and 63x 1.4 NA / Plan-Apochromat / 180 µm WD objective. Image acquisition was performed using SlideBook 6.0 (3I, Inc). 150 optical sections from five random regions of interest (ROIs) from each sample were acquired from the top of the differentiated epithelial cells. Image processing was performed using Fiji/ImageJ (Version 2.1.0/1.53c) as follows: Background was reduced using the Substract Backgound 50 pixel rolling ball radius, and the mean fluorescence intensity (MFI, a.u. arbitrary units) was measured from the average intensity images.

### Western Blotting

For total cell lysates, cells were washed twice with cold PBS and lysed with SDS/PBS lysis buffer (2% SDS/PBS buffer, 10% 10X PhosSTOP, 4% 25X protease inhibitor). Pierce BCA protein assay kit (Thermo Fisher Scientific) was used to equalize protein amounts. After adding LDS buffer (4X) with reducing agent (10X), mix protein samples were subsequently boiled at 100°C for 10 minutes to denature proteins. Proteins were separated on 4-15% mini protean TGX precast gels (Biorad) in running buffer (200 mM Glycine, 25 mM Tris, 0.1% SDS, pH8.6), transferred to nitrocellulose membrane (Cat#1620112, BioRad) in blot buffer (48 nM Tris, 39 nM Glycine, 0.04% SDS, 20% MeOH) and subsequently blocked with 5% (w/v) BSA in Tris-buffered saline with Tween 20 (TBST) for 30 minutes. The immunoblots were analyzed using primary antibodies incubated overnight at 4°C and secondary antibodies linked to horseradish peroxidase (HRP) (Invitrogen), and after each step immunoblots were washed 3X with TBST. HRP signals were visualized by enhanced chemiluminescence (ECL) (BioRad) and imaged with a AI600 Chemiluminescent Imager (Cytiva). Primary antibodies include rabbit GAPDH (14C10) monoclonal antibody (1:2500 dilution, Cat#2118, Cell Signaling Technology), rabbit anti-SARS-CoV-2 Nucleoprotein/NP antibody (1:1000 dilution, Cat#40143-R040, Sino Biological), goat polyclonal ACE2 (1:500 dilution, Cat#AF933, R&D Systems), rabbit anti-TMPRSS2 antibody (1:1000 dilution, Cat#ab109131, Abcam). ImageJ (version 1.53) was used to quantify the protein expression level relative to GAPDH levels.

### Cytokine levels

Cytokine levels were measured by AlphaLISA (PekinElmer) according to the manufacturer’s instructions.

### Quantification of infectious virus

SARS-CoV-2 titers in cell culture supernatants were determined by plaque assay on Vero cells, as described previously [28].

### RNA extraction and quantitative Reverse Transcription PCR (qRT-PCR)

RNA was extracted from NECs using Nucleozole reagent according to the manufacturer’s instructions, DNA was removed by DNase I (Thermo Fisher Scientific) treatment and 1 µg DNA-free RNA was reverse transcribed into cDNA using the High Capacity cDNA Reverse Transcription Kit (Applied Biosystems) on a Mastercycler Thermocycler (Eppendorf, Hamburg, Germany) according to the manufacturer’s instructions using random primers. Real-time PCR was performed on generated cDNA with SYBER Green (Invitrogen) using QuantStudio 6 Flex Real-Time PCR System, an Applied Biosystems Real-Time PCR Instruments (Thermo Fisher Scientific). Gene expression was normalized relative to glyceraldehyde 3-phosphate dehydrogenase (*GAPDH*) expression, fold change was calculated using the ΔΔCt method. All primers used in this study are listed in Table 1.

**Table 1.**
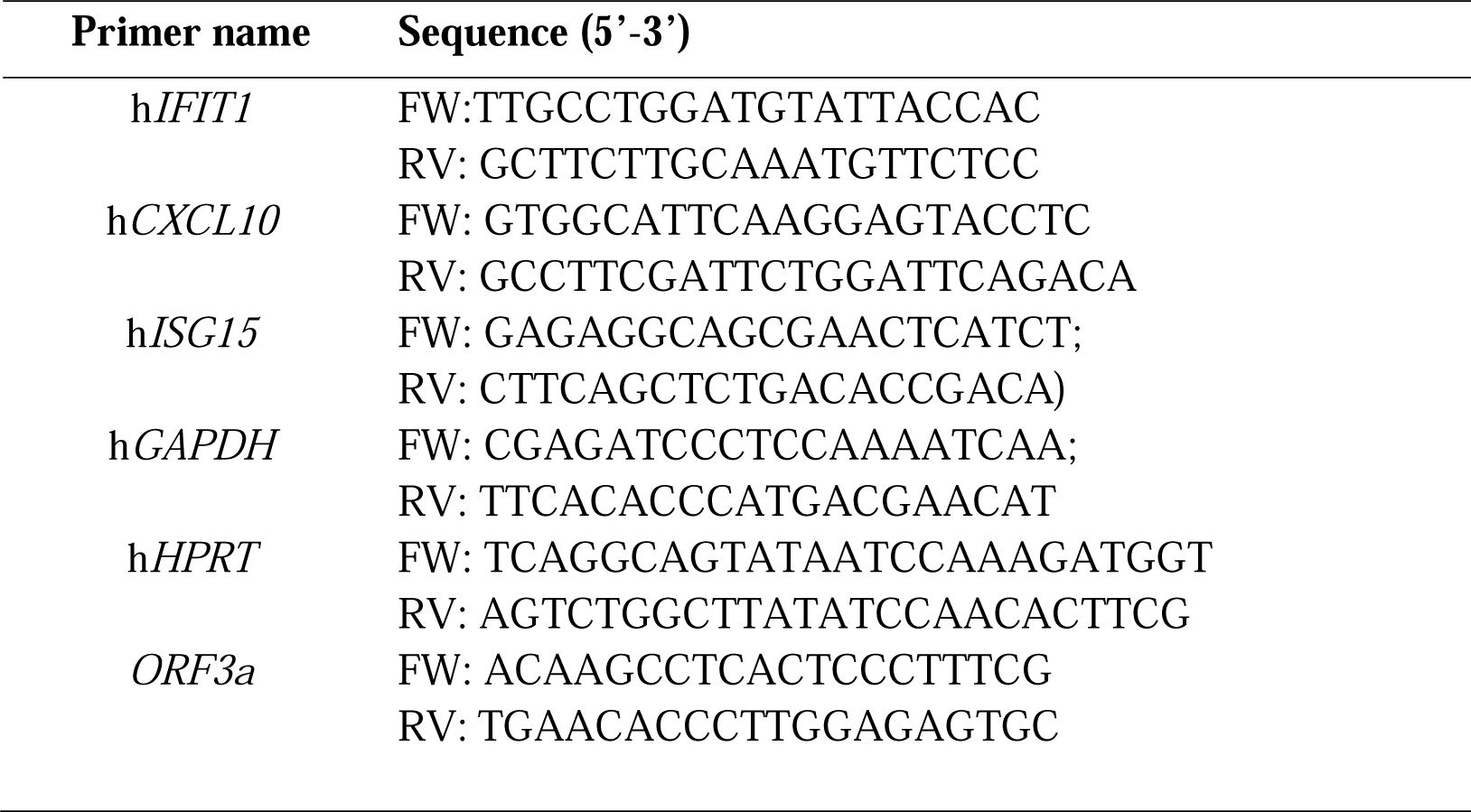
Primers used in the present study

### RNA Sequencing

RNA-Seq libraries were prepared using the Illumina stranded total RNA prep ligation with the Ribo-Zero plus kit (Illumina) and IDT for Illumina RNA UD Indexes according to the standard manufacturer’s protocol. Briefly, 50ng of total RNA was depleted of rRNA and then fragmented by heat. cDNA was synthesized from the fragmented RNA using random primers. The first strand cDNA was converted into dsDNA in the presence of dUTP to maintain the ‘strandedness’ of the library. The 3’ ends of the cDNA were adenylated and pre-index anchors were ligated. The libraries were then amplified with 14-16 cycles of PCR incorporating unique indexes for each sample to produce libraries ready for sequencing. The libraries were quantified on the Perkin Elmer LabChip GX Touch with the DNA High Sensitivity Reagent kit (Perkin Elmer). Libraries were pooled in equimolar ratios, and the pool was quantified by qPCR using the KAPA Library Quantification Kit - illumina/Universal (KAPA Biosystems) in combination with the Life Technologies Viia 7 real time PCR instrument.

Sequencing was performed using the Illumina NextSeq500 (NextSeq control software v2.2.0/Real Time Analysis v2.4.11). The library pool was diluted and denatured according to the standard NextSeq protocol and sequenced to generate single-end 76 bp reads using a 75 cycle NextSeq500/550 High Output reagent Kit v2.5 (Illumina). After sequencing, fastq files were generated using bcl2fastq2 (v2.20.0.422), which included trimming the first cycle of the insert read. Library preparation and sequencing was performed at the Institute for Molecular Bioscience Sequencing Facility (University of Queensland).

### RNA Sequencing analysis

The quality of the trimmed RNA-seq reads was assessed with FastQC [29] and MultiQC [30]. Salmon [31] was used for transcript quantification from human transcriptome (GENCODE Release 36, accessed in December 2020). A decoy aware transcriptome file was created for Salmon transcript quantification followed by the transcriptome index [31]. The R package, DESeq2 [32] was then used for differential gene expression (DGE) analysis and further validated through using the limma R package [33] with Voom transformation [34]. DGEs between virus and mock infected samples were analyzed by controlling the effect of the age group and gender of the individual samples, genes with adjusted *p*-value less than 0.05 were considered significant. Gene set enrichment analysis was performed using the R package GOseq [35]. All the R scripts were run on R-Studio platform (RStudio Team 2020, v 1.4.1717).

### Code and data availability

RNA-seq data is deposited at European Nucleotide Archive under the project– PRJEB43102. The scripts used for RNA-seq data analysis including differential gene expression and gene set enrichment analysis can be found in https://github.com/akaraw/Yanshan_Zhu_et_al.

### Statistical analysis

Where sufficient cell numbers were present, samples were performed in duplicate, and the results were averaged and shown as a single data point. If sufficient cells were not present, a single transwell was used to determine the response of that donor to viral infection. Outliers of continual variables were removed using ROUT’s test (Q = 1%). Data were tested for normality using the Shapiro-Wilk test. Where data were normally distributed, data was analyzed using an unpaired two-tailed student’s t-test. Where data were not normally distributed, data was analyzed using a Mann-Whitney U test. Significance was set at *p*<0.05. Each donor is shown with a distinct symbol that is used consistently throughout the paper.

## Results

### Pediatric nasal epithelial cells are phenotypically different to adult nasal epithelial cells

To investigate the role of NECs in SARS-CoV-2 infection, adult and pediatric NECs were differentiated at an air-liquid interface. The phenotype of these cells at baseline (i.e., prior to infection) was then assessed. Adult NECs grew as a pseudostratified columnar epithelium with scattered goblet cells and ciliated epithelial cells (Fig 1A). Pediatric NECs also grew as a pseudostratified columnar epithelium with ciliated epithelial cells and goblet cells (Fig 1A). However, scattered cells with pyknotic nucleus and condensed cytoplasm were also observed, leaving pseudocysts in the epithelium (Supplementary Figure 1). This is potentially indicative of higher cell turn-over and metabolic rate in the pediatric epithelial cells [36, 37]. Immunofluorescence images of zonal occludens-1 (ZO-1) stained NECs showed that tight junction proteins were built up closely towards the apical region of both adult and pediatric cells (Fig 1B). PAS staining indicated the presence of mucus producing cells (Fig 1A) in both pediatric and adult NECs. Consistent with these data, MUC5AC staining was detected exclusively on the apical layer, thus demonstrating mucus secretion by differentiated NECs (Fig 1B). Previous mRNA expression studies suggest that pediatric NECs express lower levels of *ACE2* and *TMPRSS2* compared to their adult counterparts [5, 9]. However, these findings are inconsistent between patient cohorts and have not been investigated at a protein level [7]. Immunofluorescence staining suggested that pediatric NECs had a trend towards lower surface levels of ACE2 compared to their adult counterparts (Fig 1B; Fig 1C). Accordingly, we sought to confirm these data using western blot on the NECs from a larger number of donors (n = 5) (Fig 1D). Whilst the same trend was observed by western blot (increased levels of ACE2 in adult NECs) this failed to reach statistical significance (Fig 1D). There was no observable trend in TMPRSS2 levels between adult and pediatric NECs (Fig 1D).

**Fig 1.**
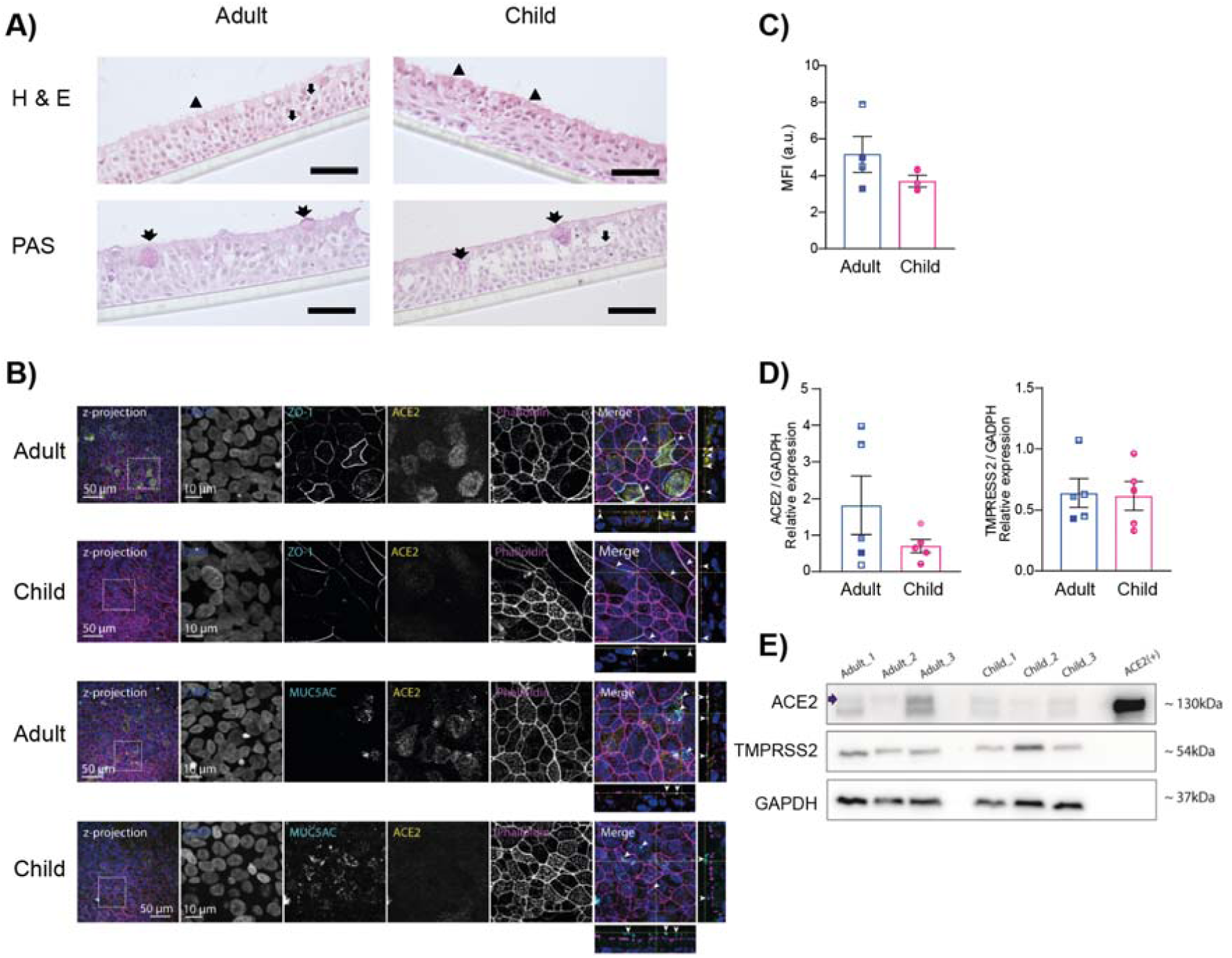
Pediatric nasal epithelial cells are phenotypically different to adult nasal epithelial cells. **A)** Representative H&E and PAS-stained sections of pediatric and adult NECs culture differentiated at an air-liquid interface (representative of 2 adults (1 female, 1 male) and 3 pediatric (1 female, 2 males) donors). Arrowheads indicate ciliated cells, arrows indicate goblet cells and double-tailed arrows indicate mucus producing cells as determined by PAS staining. Images taken at 400x magnification. Each scale bar is equivalent of 150 um **B)** Representative z-projections (150 optical sections) of pediatric and adult NECs cultures differentiated at an air-liquid interface and immunolabelled against endogenous ZO-1 and ACE2 (cyan and yellow, respectively, top panels), and MUC5AC and ACE2 (cyan and yellow, respectively, bottom panels). Cells were also stained with DAPI (blue) and phalloidin (magenta) to indicate the nucleus and actin filaments, respectively. The area in the dotted box in the images on left are shown magnified in the respective rows (10 µm bar applies to all images in the row). The merged image on right shows the orthogonal view of the z-stacks. The arrowheads indicate the zonal occludens-1 (ZO-1) stained profiles (top panels) and mucus secretion (MUC5AC) in the lower panels. **C)** Quantification of ACE2 immunofluorescence as described in the Materials and Methods. Mean ± SEM is shown. Each data point represents the average of five separate images taken from one donor (adult (N=3, 2 females, 1 male) and pediatric (N=3, 2 females, 1 male)). **D)** Relative ACE2 and TMPRSS2 protein levels compared to GAPDH in adult (3 females, 2 males) and pediatric (3 females, 2 males) NECs. Each data point represents a different donor. Mean ± SEM is shown. **E)** Representative western blot of NECs from three adult and three pediatric donors blotted for ACE2, TMPRSS2 and GAPDH. ACE2 is indicated with an arrow. Each donor is indicated by unique symbol that is used consistently throughout all figures.

### Pediatric nasal epithelial cells are less permissive to SARS-CoV-2 replication

We next sought to determine if pediatric NECs were less susceptible than adult NECs to SARS-COV-2 replication with the ancestral virus (QLD/02). Strikingly, significantly reduced SARS-CoV-2 replication was observed in pediatric NECs at 24- and 48-hours post-infection (h.p.i) (Fig 2A). Reduced SARS-CoV-2 N protein level was also observed in pediatric NECs at 24 h.p.i and 72 h.p.i (Figs 2B and 2C, Supplementary Figure 2). RNA levels showed a similar pattern although no statistical significance was recorded (Fig. 2D). After infection there was also no significant difference in ACE2 levels between adult and pediatric donors (Supplementary Figure 3).

**Fig 2.**
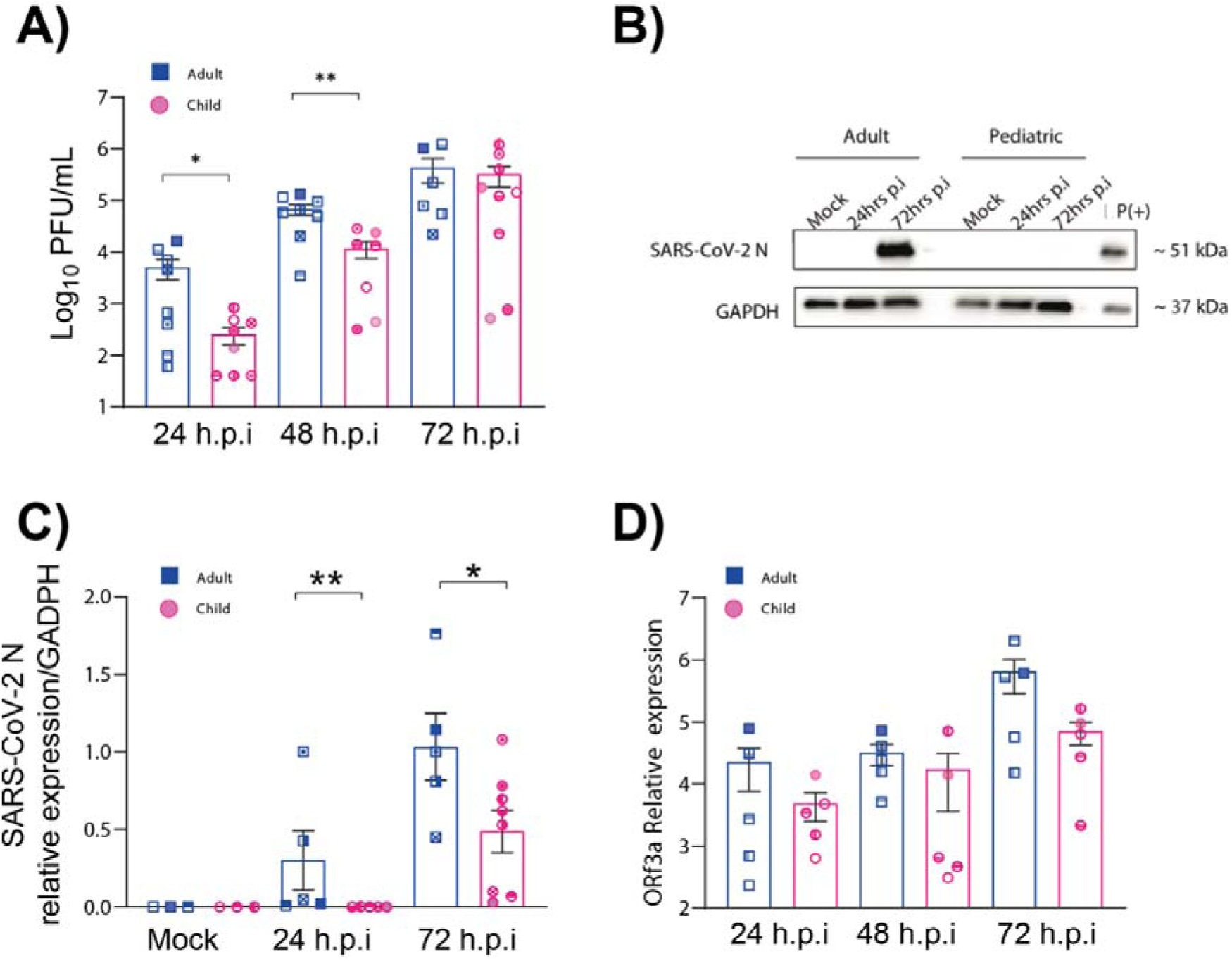
Lower replication of SARS-CoV-2 in pediatric nasal epithelial cells. **A**) Plaque forming units (PFU) of SARS-CoV-2 (QLD02) from the apical surface of nasal epithelial cells (N=8 adults: 5 females, 3 males and N=10 children: 5 females, 5 males) obtained at 24, 48 and 72 h.p.i. Cells were infected with 1.25 × 10^5^ PFU **B**) Representative western blot of adult and pediatric donor blotted for SARS-CoV-2 N at various timepoints post-infection. All the western blot results for SARS-CoV-2 N are shown as Fig. S2 **C**) Relative SARS-CoV-2 N levels compared to GAPDH in pediatric and adult NECs. **D**) Expression of ORF3a RNA in infected cells at various timepoints relative to HPRT expression. Each donor is indicated by unique symbol that is used consistently throughout all figures. Mean ± SEM is shown. *p* <0.05*, *p* <0.01**. Statistical analysis performed as described in the Materials and Methods.

### Pediatric nasal epithelial cells mount a strong anti-viral response to SARS-CoV-2

To gain a further insight into the observed decrease of SARS-CoV-2 replication in pediatric NECs RNA Seq was performed on infected adult and pediatric cells 72 hours post-infection with the ancestral SARS-CoV-2 virus. PCA analysis showed that infected cells formed distinct clusters depending on whether they were derived from pediatric or adult donors (Fig 3A). Numerous differentially expressed genes were recorded in infected cells (Fig 3B). In infected pediatric NECs, gene ontology (GO) enrichment analysis (Fig 3C) demonstrated a strong interferon response, with GO terms such as ‘viral process’, ‘type I interferon signaling’, ‘response to virus’, ‘regulation of defense response to virus’, ‘negative regulation of viral genome replication’, ‘defense response to virus’ and ‘cellular response to interferon alpha’. None of these GO terms were identified amongst the top differentially expressed GO terms in adult cells infected with SARS-CoV-2 (Fig 3D). In contrast, GO terms such as ‘cellular response to sterol’, ‘Wnt signalling pathway’ and ‘response to tumor necrosis factor’ were recorded. To confirm that these data were not restricted to a DESeq2 analysis, gene expression data were also analyzed using limma (Supplementary Table 1 and 2). Once again, in infected pediatric NECs GO terms such as ‘response to virus’, ‘cellular response to cytokine stimulus’ and ‘defense response to virus’ were recorded. In contrast, infected adult NECs were associated with GO terms such as ‘detection of stimulus involved in sensory perception’ and ‘sensory perception’. To further validate these data, we assessed gene expression by qPCR of three genes associated with inflammatory/anti-viral response - interferon-induced protein with tetratricopeptide repeats 1 (*IFIT1*); C-X-C motif chemokine ligand 10 (*CXCL10*) and interferon stimulated gene 15 (*ISG15*). Infected pediatric NECs had significantly higher levels of *IFIT1* compared to infected adult NECs (Fig 4A). Infected pediatric NECs also had a trend of increased IFN-alpha, IFN-beta and CXCL10 protein levels following SARS-CoV-2 infection, although donor-to-donor variability precluded significance (Fig. 4B-D).

**Fig 3.**
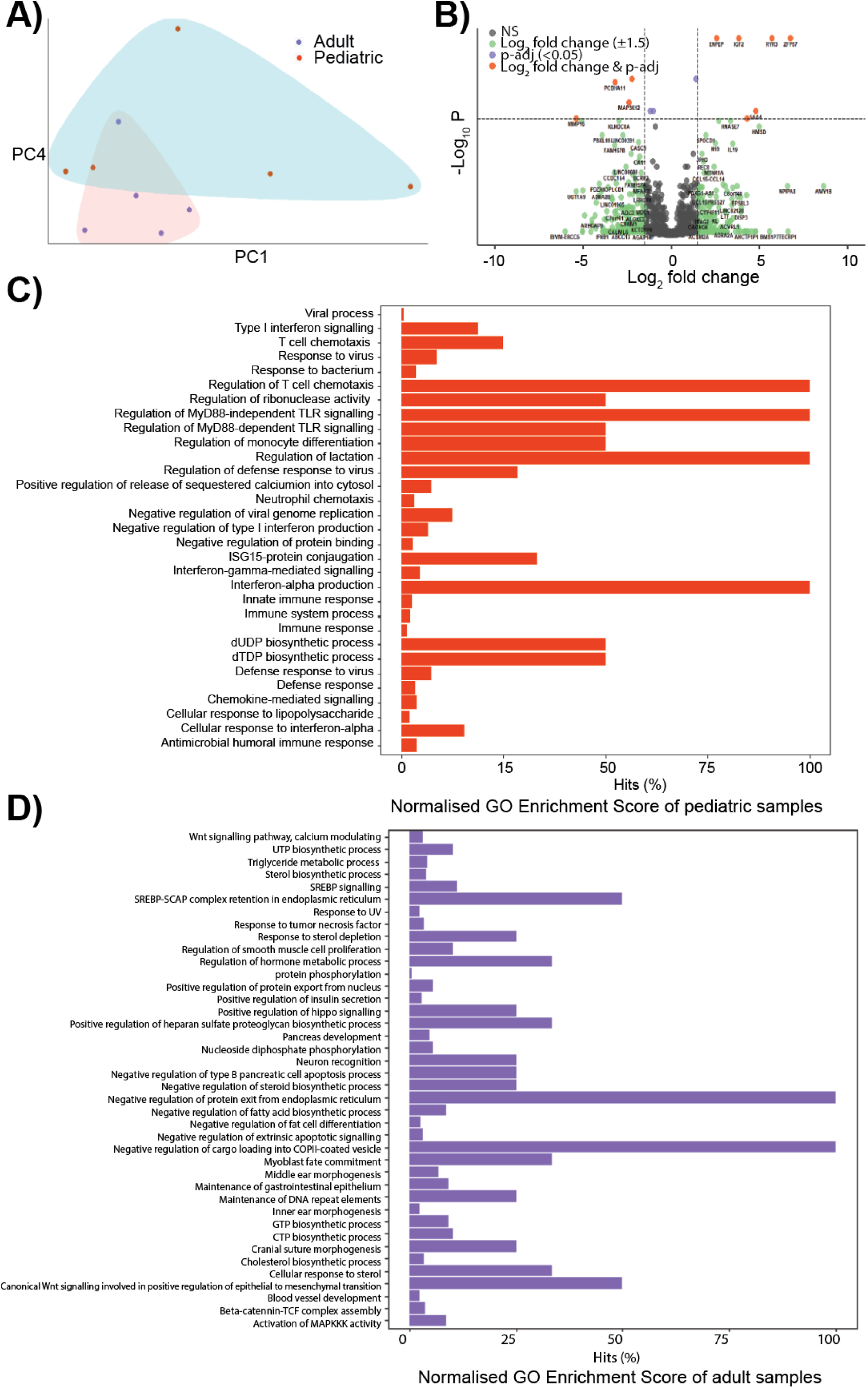
Pediatric epithelial cells have a different transcriptional response to SARS-CoV-2. **A)** Principal component analysis for the global transcriptional response of naive pediatric and adult NECs. Data points represent individual donors (N=5 adults: 4 females, 1 male and N = 5 pediatric donors: 2 females, 3 males). **B)** Volcano plot illustrating differentially expressed genes (DEGs) of infected pediatric NECs compared to adult cells. DEGs statistically different between the two patient groups with a fold change of >1.5 are indicated in orange. DEGs statistically different between two groups with a fold change of <1.5 are shown in purple. DEGs not statistically different between two groups with a fold change of >1.5 are shown in green. NS = not significant. **C**) Gene ontology (GO) analysis of DEGs in infected pediatric NECs were displayed by the bar chart. The bars of significantly GO enriched (Overrepresented p value < 0.05) results were marked in red, x-axis reflects the gene count hits as a percentage over genes in each GO category; y-axis reflects different GO terms. **D)** Gene ontology (GO) analysis of DEGs in adult NECs were displayed by the bar chart. The bars of significantly enriched GO (Overrepresented p value < 0.05) enrichment results were marked in purple and represents the gene count hits (as a percentage over number of genes in a given category); y-axis reflects different GO terms.

**Fig 4.**
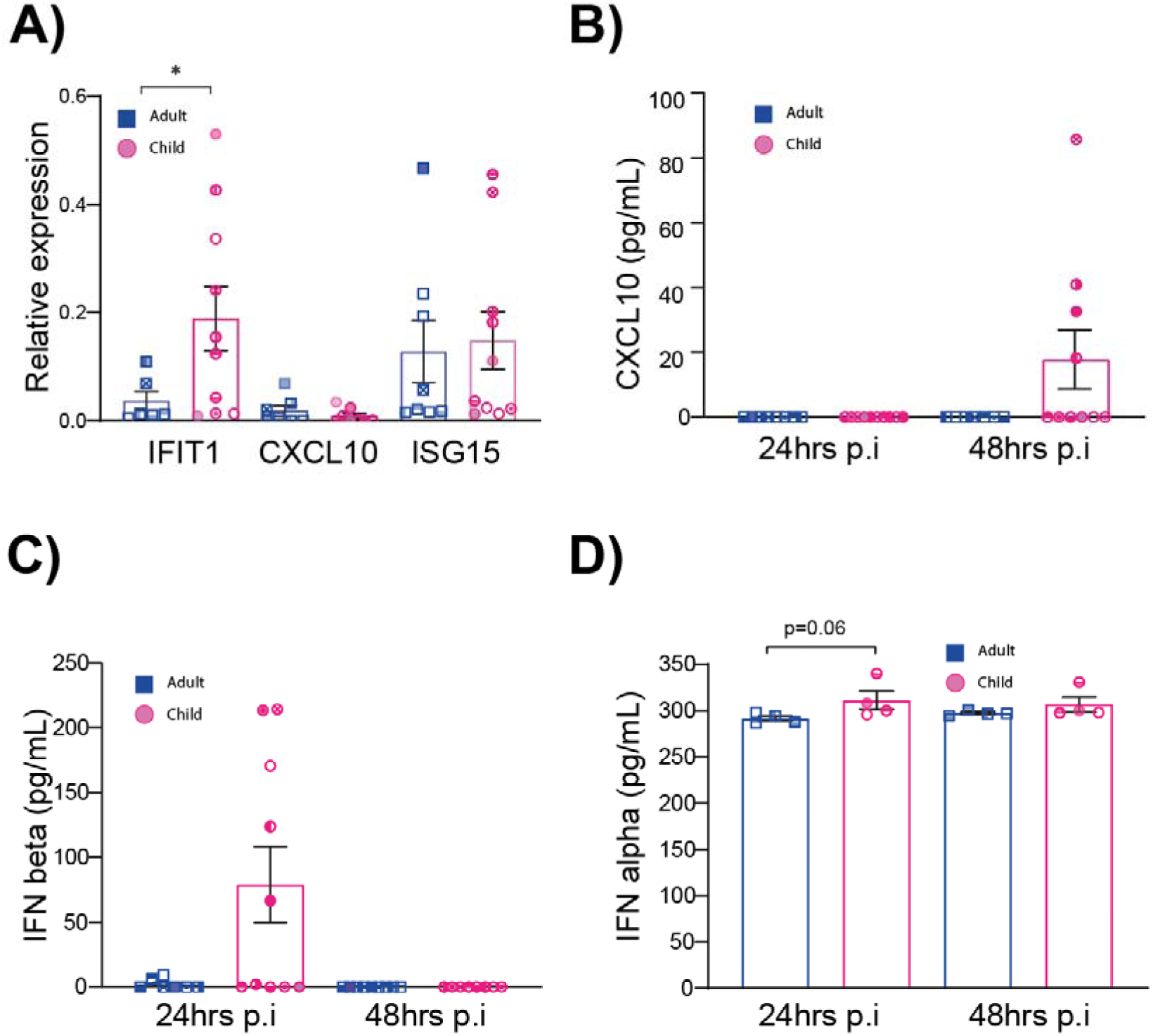
Pediatric epithelial cells have a stronger type I IFN response to viral infection. **A)** Expression of interferon associated genes in ancestral SARS-CoV-2 (QLD02) infected epithelial cells relative to uninfected controls (48 hours post-infection). Each data point represents a different donor (N=8 adults: 5 females, 3 males and N=10 pediatric: 5 females, 5 males). Gene expression (fold change) was calculated using the ΔΔCt method relative to GAPDH expression. **B-D)** Levels of IFN-alpha, IFN-beta and CXCL10 in epithelial cell supernatant at three timepoints post-infection (p.i) with ancestral SARS-CoV-2 (QLD02). Mean ± SEM is shown. *p* <0.05*, statistical analysis performed as described in the Materials and Methods. Each donor is indicated by unique symbol that is used consistently throughout all figures. *Number of donors shown in 4B&C (N=8 adults: 5 females, 3 males and N=10 children: 5 females, 5 males) and 4D (N=5 adults: 4 females, 1 male and N=5 children: 2 females, 3 males) were different.

We next sought to investigate if we observed similar phenotype in pediatric epithelial cells infected with selected variants of concern (Delta, Omicron). At 24 hours post-infection there were significantly high titres of infectious virus and viral RNA in adult epithelial cells infected with the ancestral virus (QLD02) and Delta compared to the epithelial cells of children (Figure 5). Interestingly, whilst a similar (albeit not significant) trend was observed in Omicron infectious virus titres there was no difference in Omicron RNA levels in pediatric vs adult NECs (Figure 5).

**Fig 5.**
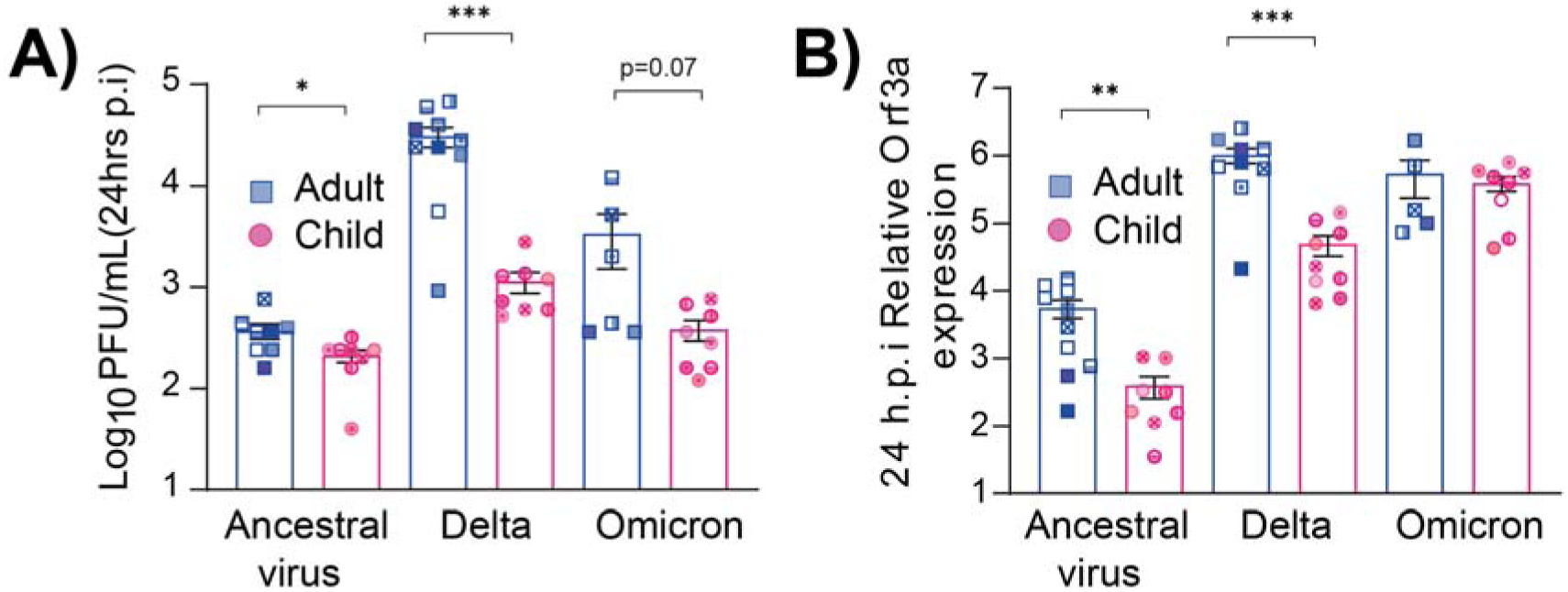
Lower replication of SARS-CoV-2 Delta in pediatric nasal epithelial cells. **A**) Plaque forming units (PFU) of SARS-CoV-2 from the apical surface of nasal epithelial cells NECs obtained at 24 hrs post-infection (p.i). Cells were infected with 1.7 × 10^4^ PFU of each of the respective viral variants. **B)** Expression of ORF3a RNA in infected cells at relative to GAPDH expression. Each donor is indicated by unique symbol that is used consistently throughout all figures (N=10 adults: 5 females, 5 males and N=10 children: 5 females, 5 males). Mean ± SEM is shown. *p* <0.05*, *p* <0.01**. Statistical analysis performed as described in the Materials and Methods.

## Discussion

Large clinical data sets and systematic reviews suggest that children are less often infected with the ancestral SARS-CoV-2 and have less severe symptoms than adults [38-41]. However, the mechanisms driving these observations have been unclear. Here, we have provided the first experimental evidence that the pediatric nasal epithelium may play an important role in reducing the susceptibility of children to SARS-CoV-2.

Previous studies have suggested that the reduced susceptibility of children to SARS-CoV-2 infection is due to reduced mRNA expression of SARS-CoV-2 receptors ACE2 and TMPRSS2. Specifically, it has been proposed that the lower level of ACE2 and TMPRSS2 in pediatric upper airways epithelial cells limits disease severity and viral infectivity in children [5, 42], although this has remained somewhat controversial [6, 7]. In the present study, whilst there was a trend towards decreased ACE2 protein levels in pediatric NECs there was significant donor-to-donor variability that precluded statistical significance. We interpret these data as suggesting that ACE2 levels may contribute to, but are not the sole factor, in the increased resistance of children to SARS-CoV-2.

Despite donor-to-donor differences in ACE2 expression, we consistently observed a significant reduction in ancestral SARS-CoV-2 (QLD02) replication in pediatric NECs compared to NECs of adults. Given that the nasal epithelium is the first site of SARS-CoV-2 infection these data are consistent with the reduced number of SARS-CoV-2 infected children recorded in household and school transmission studies [43, 44]. However, we recognise it is challenging to compare data from controlled experimental studies to data obtained from patient sampling, where it is difficult to control for time of sampling relative to the onset of infection. Rather, decreased viral replication in pediatric epithelial cells is consistent with experimental studies in ferrets where aged ferrets showed higher viral load and longer nasal virus shedding [45].

Consistent with reduced ancestral SARS-CoV-2 replication in the nasal epithelium of children, pediatric epithelial cells had a more pronounced pro-inflammatory response (compared to adult cells) following an ancestral SARS-CoV-2 infection. In particular, a pronounced interferon response and the expression of interferon stimulated genes (*ISGs*) was higher in infected pediatric, compared to adult, NECs. Increased *ISG* expression, and the subsequent anti-viral response may contribute to the reduced viral replication observed in pediatric cells. Importantly, unlike the lower respiratory tract, any resultant cell death or immunopathology in the upper respiratory tract is unlikely to lead to respiratory distress and therefore remains beneficial to the host [12]. These findings are consistent with those of Maughan et al, who analyzed transcriptional profile of airway (tracheobronchial) epithelium and observed upregulated type I and II *IFNs* associated genes in children [10]. Similarly, in the nasal fluid of children and adults presenting to the emergency department with SARS-CoV-2 there were significantly higher levels of *IFN-*α*2* in the fluid derived from children. Increased interferon signaling was also recorded in the nasopharyngeal transcriptome of children compared to that of adults during early SARS-CoV-2 infection [6, 46].

The question remains as to why pediatric epithelial cells mount a stronger inflammatory and anti-viral response to ancestral SARS-CoV-2 compared to adult cells. This may represent an adaptation to the increased antigenic challenge observed in childhood. Alternatively, it is possible that increased antigenic exposure in childhood ‘trains’ nasal epithelium in children to mount a stronger pro-inflammatory response to any antigenic challenge. It is also possible that metabolic differences between pediatric and NECs (as potentially suggested by the different morphologies of the cells) could alter gene expression.

It is striking to note that the VOC Delta also replicated significantly better in the NECs of adults compared to those of children. These data suggest that any increase in pediatric infections during the Delta wave are unlikely to be due to the fact that the virus has substantially evaded the innate immune response of pediatric NECs and are instead more likely attributable to other factors (e.g. age dependent differences in prior infection and/or vaccination). Interestingly, the Delta variant replicated to higher titres in the NECs of both adults and children when compared to the ancestral virus. This is consistent with previous studies [47] and may be associated with the increased transmissibility of Delta compared to the ancestral virus. Interestingly, at least in adults, the more recent Omicron variant did not replicate more efficiently in the NECs than the earlier Delta variant. These data support suggestions that Omicron does not necessarily have a replicative advantage over Delta in the URT of adults [48], and the observed increase in transmission with Omicron in adults is more likely reflective of increased antibody evasion [49-52].

Surprisingly, age-dependent differences in viral replication in NECs was much less pronounced in the case of Omicron. Indeed, there was no notable difference in Omicron RNA titres at 24 hours post-infection in adult vs pediatric NECs. These data may provide preliminary evidence that the Omicron variant, at least to some extent, is able to evade aspects of the pediatric innate immune response, as has previously been demonstrated [53]. Whether this is sufficient to result in an increased number of pediatric infections during the Omicron wave [54, 55], or whether other factors are more important, remains to be determined. However, it is striking to note (at least in terms of RNA levels) increased titres of Omicron in pediatric NECs compared to infection with Delta and the ancestral virus. These data are consistent with the increased number of pediatric infections observed during the Omicron wave [54, 55].

Finally, it is important to recognize the limitations of this study. Due to the difficulties associated with obtaining NECs from children only a limited number of donors could be used for this study. However, as donors were not selected according to susceptibility to respiratory viral infection, their responses should be broadly representative of healthy children. Furthermore, our data focused on the role of nasal epithelial cells in age-dependent differences in SARS-CoV-2 infection. However, there may be other mechanisms to explain the reduced susceptibility of children to SARS-CoV-2 infection that were not measured in the present study. For example, children and adolescents have higher titers of preexisting antibodies to SARS-CoV-2 compared to adults [56]. This study is unable to ascertain if this plays a more significant role than the nasal epithelium in protecting children from infection *in vivo*.

## Conclusions

The data presented here strongly suggest that the nasal epithelium of children is distinct and that it may afford children some level of protection from ancestral SARS-CoV-2, although such age-dependent differences become less pronounced in the case of Omicron infection.

## Acknowledgments

We greatly thank the participants in the study and the members of the research team. The authors thank the participants and their families in the Western Australian Epithelial Research Program (WAERP) for their contribution to this study. This program has in part been funded by the Western Australian Future Health Research & Innovation Fund. Current members of WAERP include; Anthony Kicic, Stephen M. Stick, Elizabeth Kicic-Starcevich, Amy Greenly, Angela Fuery, Luke W. Garratt, Erika N. Sutanto, Kevin Looi, Jessica Hillas, Thomas Iosifidis, Craig Schofield, Samantha McLean, Luke Berry, Samuel T. Montgomery, Kak-Ming Ling, Renee Ng, Andrew Vaitekenas, Daniel Laucirica, Matthew Poh, Reanne Ho, Joshua Iszatt, Katherine Landwehr, Denby Evans, Rebecca Watkinson, Patricia Agudelo-Romero, Jose Caparros-Martin, Alexander de Bont, Julia Maynard, Rohan Flint, Jack Canning, Shyan Vijayasekaran, George Sim, Mairead Heaney, Tom Rawlings, Neil Chambers, Christopher Johnson, Eugene Roscioli, Alexander Larcombe, Tim Barnett, Rael, Rivers, Kate McGee.

## Supporting information

**Supplementary Figure 1:**
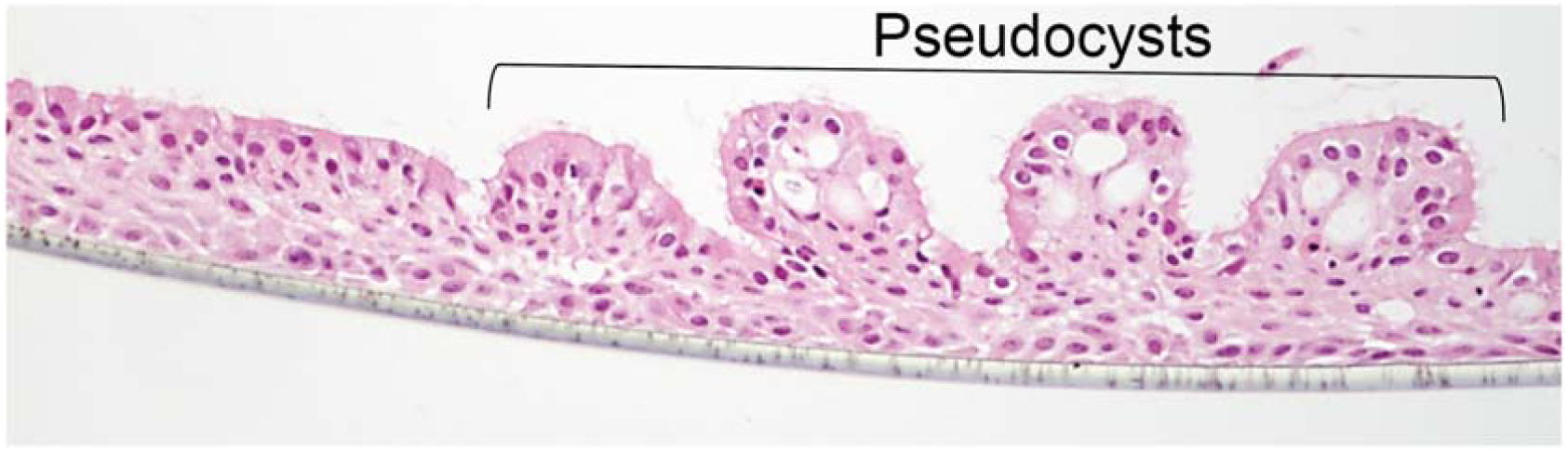
Pediatric nasal epithelial cells display pseudocysts. Representative Hematoxylin & Eosin-stained section of pediatric NECs culture differentiated at an air-liquid interface.

**Supplementary Figure 2:**
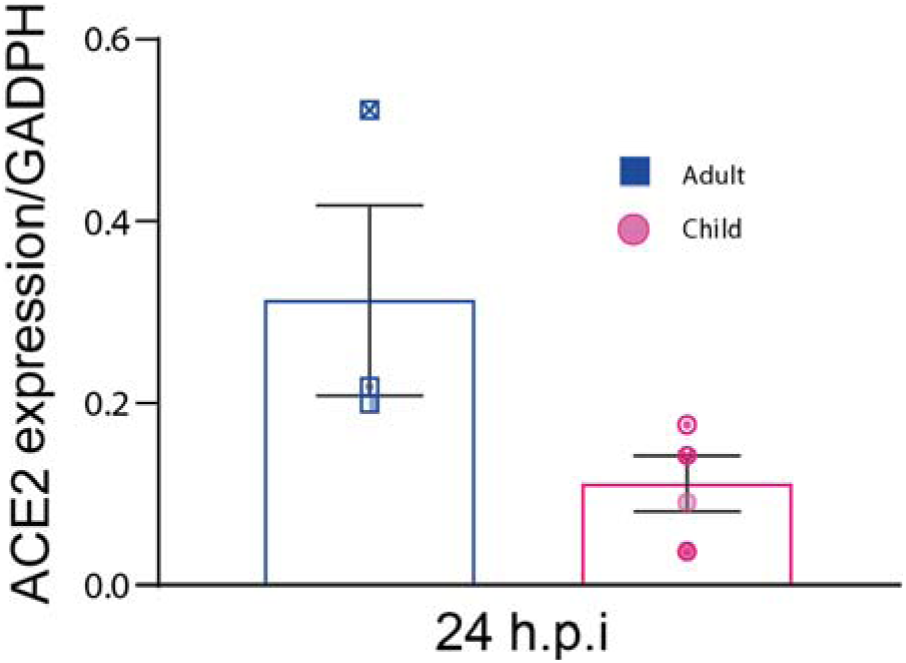
Relative SARS-CoV-2 NP levels compared to GAPDH in pediatric and adult NECs. Western blot of adults and pediatric donors blotted for SARS-CoV-2 N at 24 hours post-infection (N=3 adults (1 females, 2 male) and N = 5 children (3 female, 2 males)).

**Supplementary Fig. 3:**
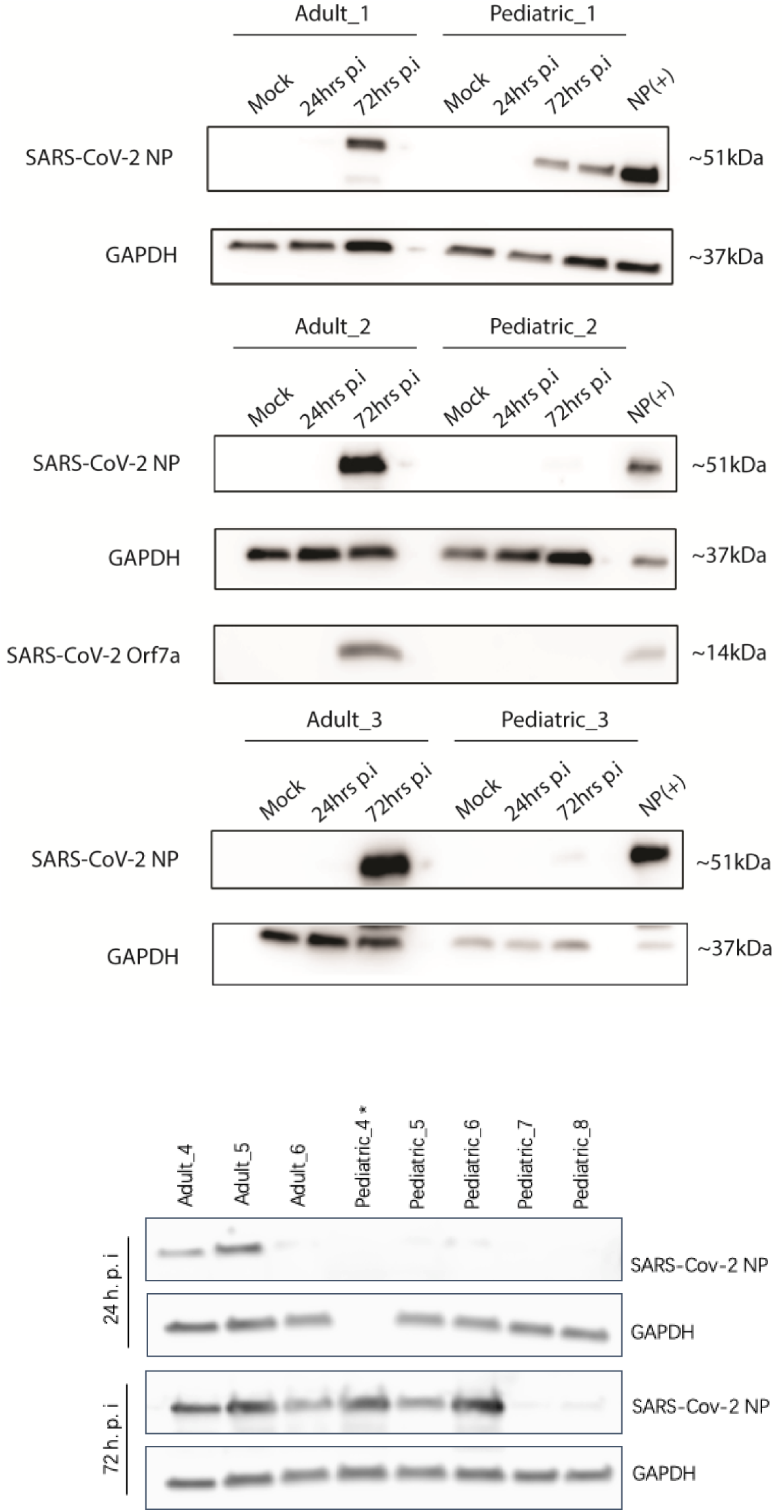
Relative ACE2 levels compared to GAPDH in pediatric and adult NECs. Western blot of adults and pediatric donors blotted for ACE at various timepoints post-infection (N= 6 adults (3 females, 3 male) and N = 10 children (5 female, 5 males)).

