## Supplemental tables for "Pediatric nasal epithelial cells are less permissive to SARS-CoV-2 replication compared to adult cells"

**Table E1. Differentially expressed genes (DEGs) of infected pediatric NECs (72 hours post-infection) compared to adult cells.** DEGs were identified using DESeq2, genes with adjusted p-value less than 0.05 value were considered significant.

| Genes | **baseMean** | **log2FC** | **lfcSE** | **stat** | **pvalue** | **padj** |
| --- | --- | --- | --- | --- | --- | --- |
| ADH7 | 2874.532 | -1.00421 | 0.237966 | -4.21997 | 2.44E-05 | 0.041236 |
| ENPEP | 32.35705 | 2.58263 | 0.525359 | 4.915929 | 8.84E-07 | 0.006328 |
| GPATCH4 | 51.0056 | -1.19709 | 0.284169 | -4.21259 | 2.52E-05 | 0.041236 |
| IGF2 | 24.55162 | 3.851189 | 0.793619 | 4.852693 | 1.22E-06 | 0.006328 |
| MAP3K12 | 151.4382 | -2.38054 | 0.549252 | -4.33415 | 1.46E-05 | 0.032863 |
| MMP16 | 4.66984 | -5.37427 | 1.300314 | -4.13306 | 3.58E-05 | 0.049609 |
| PCDHA11 | 14.09029 | -3.17871 | 0.710306 | -4.47513 | 7.64E-06 | 0.019601 |
| PTPRQ | 60.26947 | -2.23112 | 0.492802 | -4.52742 | 5.97E-06 | 0.017879 |
| RYR3 | 99.46414 | 5.719201 | 1.185634 | 4.82375 | 1.41E-06 | 0.006328 |
| SAA4 | 8.002679 | 4.804457 | 1.13706 | 4.225334 | 2.39E-05 | 0.041236 |
| SLC26A4 | 114.124 | 4.29726 | 1.039886 | 4.132433 | 3.59E-05 | 0.049609 |
| ZBED6CL | 109.8951 | 1.398693 | 0.308122 | 4.539415 | 5.64E-06 | 0.017879 |
| ZFP57 | 16.79348 | 6.78271 | 1.351402 | 5.019018 | 5.19E-07 | 0.006328 |

**Table E2. Differentially expressed genes (DEGs) of infected pediatric NECs (72 hours post-infection) compared to adult cells.** DEGs were identified using limma, genes with adjusted p-value less than 0.05 value were considered significant.

| **Genes** | **Log2FC** | **AveExpr** | **t** | **pvalue** | **padj** | **baseMean** |
| --- | --- | --- | --- | --- | --- | --- |
| TMPRSS11B | 5.25370 | 0.80005 | 12.52257 | 6.90E-09 | 0.000115039 | 8.26292 |
| MAL | 3.05471 | 1.85718 | 9.27935 | 2.99E-07 | 0.002495525 | 7.06046 |
| CCT6B | 1.79261 | 3.51197 | 8.15790 | 1.26E-06 | 0.003576789 | 5.77689 |
| XKR9 | 2.33378 | 2.89458 | 8.15097 | 1.27E-06 | 0.003576789 | 5.56156 |
| COL8A1 | 1.91266 | 3.80711 | 7.92954 | 1.75E-06 | 0.003576789 | 5.46652 |
| SPAG16-DT | 1.95120 | 3.08596 | 7.91267 | 1.79E-06 | 0.003576789 | 5.39066 |
| LINC02021 | 2.27798 | 1.61930 | 7.83067 | 2.01E-06 | 0.003576789 | 5.05908 |
| LINC01684 | 2.01391 | 3.50171 | 7.82943 | 2.02E-06 | 0.003576789 | 5.31520 |
| N4BP2L2-IT2 | 2.40897 | 3.02725 | 7.82547 | 2.07E-06 | 0.003576789 | 5.25461 |
| ROR1-AS1 | 2.13206 | 3.51829 | 7.79910 | 2.15E-06 | 0.003576789 | 5.25900 |
| RAD51-AS1 | 2.01955 | 2.67972 | 7.45260 | 3.52E-06 | 0.004483567 | 4.76776 |
| LINC02822 | 2.03322 | 3.42002 | 7.41232 | 3.84E-06 | 0.004483567 | 4.70438 |
| POU5F2 | 2.31396 | 2.97916 | 7.35883 | 4.27E-06 | 0.004483567 | 4.59467 |
| SEPSECS-AS1 | 1.56623 | 3.98177 | 7.30840 | 4.37E-06 | 0.004483567 | 4.56211 |
| TROAP | 2.32251 | 0.95461 | 7.27362 | 4.61E-06 | 0.004483567 | 4.17078 |
| TPTEP2 | 2.31685 | 2.15914 | 7.26564 | 4.66E-06 | 0.004483567 | 4.26420 |
| RRN3P3 | 1.71693 | 2.93552 | 7.21137 | 5.07E-06 | 0.004483567 | 4.42481 |
| LINC02614 | 1.86022 | 3.22350 | 7.21124 | 5.07E-06 | 0.004483567 | 4.43491 |
| PRANCR | 1.83121 | 2.90179 | 7.11293 | 5.89E-06 | 0.004483567 | 4.28234 |
| CAP2P1 | 2.92575 | 0.53164 | 7.07843 | 6.21E-06 | 0.004483567 | 3.64081 |
| FAM13A-AS1 | 2.12249 | 2.25576 | 7.05239 | 6.47E-06 | 0.004483567 | 4.14252 |
| PMS2P3 | 1.88699 | 2.19333 | 7.04357 | 6.56E-06 | 0.004483567 | 4.11961 |
| USP32P3 | 1.61880 | 3.32649 | 7.03498 | 6.65E-06 | 0.004483567 | 4.16966 |
| LINC00598 | 1.74060 | 2.87596 | 7.02714 | 6.73E-06 | 0.004483567 | 4.16146 |
| CFAP298-TCP10L | 1.65886 | 3.44009 | 7.01515 | 6.86E-06 | 0.004483567 | 4.13447 |
| PAXBP1-AS1 | 1.84730 | 3.21746 | 7.00241 | 6.99E-06 | 0.004483567 | 4.12499 |
| SILC1 | 1.85528 | 2.96720 | 6.95291 | 7.55E-06 | 0.004528195 | 4.02685 |
| ZNF519 | 1.65716 | 3.46061 | 6.94854 | 7.61E-06 | 0.004528195 | 4.03715 |
| POC1B-AS1 | 1.57505 | 3.46278 | 6.88362 | 8.42E-06 | 0.004576337 | 3.94357 |
| CATSPER2 | 2.36195 | 2.64723 | 6.87472 | 8.54E-06 | 0.004576337 | 3.83809 |
| OR9H1P | 1.77655 | 2.34875 | 6.87348 | 8.56E-06 | 0.004576337 | 3.90454 |
| LINC01176 | 2.16049 | 2.17106 | 6.85161 | 8.86E-06 | 0.004576337 | 3.86765 |
| KIN | 1.32494 | 3.77307 | 6.81971 | 9.31E-06 | 0.004576337 | 3.83454 |
| LINC02605 | 1.90571 | 3.15036 | 6.81470 | 9.39E-06 | 0.004576337 | 3.84119 |
| OR5A1 | 1.81623 | 2.46638 | 6.77308 | 1.00E-05 | 0.004576337 | 3.75773 |
| COLQ | 2.11177 | 1.28520 | 6.77120 | 1.01E-05 | 0.004576337 | 3.52134 |
| LINC02558 | 1.88799 | 2.28335 | 6.76492 | 1.02E-05 | 0.004576337 | 3.68056 |
| LINC01609 | 1.54118 | 3.81371 | 6.72804 | 1.08E-05 | 0.004580988 | 3.67859 |
| TPTEP1 | 1.70305 | 2.22455 | 6.69179 | 1.14E-05 | 0.004580988 | 3.61857 |
| DICER1-AS1 | 1.83443 | 2.56186 | 6.69004 | 1.14E-05 | 0.004580988 | 3.64577 |
| CCDC122 | 1.41614 | 3.81426 | 6.68597 | 1.15E-05 | 0.004580988 | 3.63369 |
| GHET1 | 1.95387 | 2.12620 | 6.67693 | 1.17E-05 | 0.004580988 | 3.56009 |
| LINC02018 | 1.47416 | 3.74734 | 6.67001 | 1.18E-05 | 0.004580988 | 3.58669 |
| LINC01088 | 1.65066 | 3.29809 | 6.65152 | 1.22E-05 | 0.00461148 | 3.58949 |
| LINC01422 | 2.01452 | 2.45057 | 6.59557 | 1.33E-05 | 0.004794162 | 3.46987 |
| THRB-IT1 | 2.22380 | 2.00458 | 6.57312 | 1.38E-05 | 0.004794162 | 3.33633 |
| C1RL-AS1 | 1.96295 | 3.26718 | 6.57698 | 1.43E-05 | 0.004794162 | 3.42061 |
| LINC02742 | 2.10621 | 1.82701 | 6.54280 | 1.45E-05 | 0.004794162 | 3.30899 |
| NDUFAF2 | 1.74411 | 3.42213 | 6.55465 | 1.46E-05 | 0.004794162 | 3.40574 |
| DLEU2L | 1.86114 | 1.56345 | 6.51861 | 1.51E-05 | 0.004794162 | 3.25465 |
| SAP30L-AS1 | 1.89929 | 1.84375 | 6.50514 | 1.54E-05 | 0.004794162 | 3.31839 |
| LINC02280 | 2.11415 | 1.81614 | 6.49767 | 1.56E-05 | 0.004794162 | 3.31633 |
| DPH6 | 1.69679 | 3.37609 | 6.49122 | 1.58E-05 | 0.004794162 | 3.34105 |
| LINC02324 | 2.20859 | 1.83062 | 6.48346 | 1.60E-05 | 0.004794162 | 3.23372 |
| MIATNB | 1.70885 | 2.43590 | 6.45645 | 1.67E-05 | 0.004794162 | 3.28489 |
| ACTR3C | 1.50338 | 4.30525 | 6.45998 | 1.68E-05 | 0.004794162 | 3.23849 |
| ST3GAL3 | 1.67063 | 3.77064 | 6.47306 | 1.70E-05 | 0.004794162 | 3.23312 |
| CENATAC | 1.91373 | 3.20842 | 6.47302 | 1.70E-05 | 0.004794162 | 3.27352 |
| OR4K17 | 1.97526 | 2.08009 | 6.44113 | 1.71E-05 | 0.004794162 | 3.23433 |
| LINC00271 | 1.62805 | 3.81367 | 6.43765 | 1.73E-05 | 0.004794162 | 3.23885 |
| PRNCR1 | 1.66412 | 3.96511 | 6.43685 | 1.80E-05 | 0.004794783 | 3.18851 |
| OR10K1 | 2.12868 | 3.96268 | 6.43542 | 1.80E-05 | 0.004794783 | 3.19154 |
| ZNRD1ASP | 1.41401 | 3.82191 | 6.40587 | 1.81E-05 | 0.004794783 | 3.15038 |
| CYP1B1-AS1 | 1.87269 | 2.43666 | 6.38557 | 1.87E-05 | 0.004879507 | 3.14668 |
| FAM229A | 2.68298 | 1.04513 | 6.36717 | 1.93E-05 | 0.004897532 | 2.86622 |
| CCDC144B | 1.62653 | 3.26662 | 6.36355 | 1.99E-05 | 0.004897532 | 3.09067 |
| GTF2IP20 | 1.39844 | 3.87699 | 6.34646 | 2.00E-05 | 0.004897532 | 3.04939 |
| POT1-AS1 | 1.97720 | 1.60837 | 6.34061 | 2.02E-05 | 0.004897532 | 2.96327 |
| OR10G3 | 1.96947 | 1.83397 | 6.33753 | 2.03E-05 | 0.004897532 | 3.01978 |
| SPATA6L | 1.33780 | 4.92812 | 6.31396 | 2.20E-05 | 0.005088259 | 2.88306 |
| ZNF788P | 2.23180 | 2.76832 | 6.29583 | 2.24E-05 | 0.005088259 | 3.00357 |
| CLEC2D | 1.95376 | 3.31708 | 6.29583 | 2.26E-05 | 0.005088259 | 2.99501 |
| UBE2Q2P2 | 2.26264 | 2.46271 | 6.28413 | 2.31E-05 | 0.005088259 | 2.98499 |
| GOLGA6L9 | 1.76041 | 4.35180 | 6.28192 | 2.31E-05 | 0.005088259 | 2.86619 |
| ZSCAN23 | 2.34063 | 1.52294 | 6.25314 | 2.33E-05 | 0.005088259 | 2.81922 |
| LINC01572 | 1.79418 | 2.79994 | 6.24741 | 2.35E-05 | 0.005088259 | 2.96037 |
| LINC01934 | 1.65389 | 3.77884 | 6.26566 | 2.36E-05 | 0.005088259 | 2.92867 |
| WHAMMP2 | 1.71845 | 2.32358 | 6.23766 | 2.39E-05 | 0.005088259 | 2.93732 |
| EFCAB6-AS1 | 1.86743 | 2.90774 | 6.22425 | 2.44E-05 | 0.005088259 | 2.91967 |
| LINC01224 | 1.87758 | 1.55330 | 6.22356 | 2.45E-05 | 0.005088259 | 2.88501 |
| C3orf35 | 2.09041 | 2.52783 | 6.22020 | 2.50E-05 | 0.005088259 | 2.90240 |
| UGT2A2 | 2.49019 | 3.18855 | 6.22825 | 2.53E-05 | 0.005088259 | 2.89065 |
| KCNMB2-AS1 | 1.56563 | 2.84884 | 6.18632 | 2.60E-05 | 0.005088259 | 2.86374 |
| TAMM41 | 1.68543 | 3.09960 | 6.18588 | 2.61E-05 | 0.005088259 | 2.86426 |
| CMSS1 | 1.77536 | 2.93175 | 6.16774 | 2.69E-05 | 0.005088259 | 2.83523 |
| LINC01091 | 1.30058 | 3.74579 | 6.15666 | 2.74E-05 | 0.005088259 | 2.76666 |
| LINC00882 | 1.62904 | 3.68438 | 6.17170 | 2.77E-05 | 0.005088259 | 2.76955 |
| OR2M3 | 2.09329 | 2.97690 | 6.15883 | 2.84E-05 | 0.005088259 | 2.78849 |
| LINC01001 | 1.74622 | 3.81843 | 6.15780 | 2.84E-05 | 0.005088259 | 2.73595 |
| TIGD1 | 1.61132 | 4.06871 | 6.14548 | 2.90E-05 | 0.005088259 | 2.66740 |
| RABGAP1L-IT1 | 1.85696 | 2.87227 | 6.12754 | 2.92E-05 | 0.005088259 | 2.75540 |
| C6orf163 | 1.72646 | 1.91223 | 6.10780 | 2.97E-05 | 0.005088259 | 2.72997 |
| DARS-AS1 | 1.89003 | 2.61316 | 6.10578 | 2.98E-05 | 0.005088259 | 2.73383 |
| ARL17B | 1.61845 | 3.74954 | 6.10532 | 2.98E-05 | 0.005088259 | 2.71877 |
| LINC00891 | 1.82246 | 2.87973 | 6.10423 | 2.99E-05 | 0.005088259 | 2.73463 |
| ACYP1 | 1.54717 | 2.54852 | 6.09909 | 3.01E-05 | 0.005088259 | 2.72417 |
| ZNF235 | 1.37443 | 3.08838 | 6.09344 | 3.04E-05 | 0.005088259 | 2.70863 |
| FAM215B | 2.22099 | 0.92166 | 6.08139 | 3.10E-05 | 0.005088259 | 2.63594 |
| HYKK | 1.33920 | 3.65982 | 6.07992 | 3.11E-05 | 0.005088259 | 2.67039 |
| ABCC9 | 1.59000 | 3.75506 | 6.10278 | 3.11E-05 | 0.005088259 | 2.61802 |
| LINC02080 | 1.56288 | 2.07974 | 6.07322 | 3.15E-05 | 0.005088259 | 2.68022 |
| TMEM9B-AS1 | 1.61346 | 3.41854 | 6.09290 | 3.15E-05 | 0.005088259 | 2.64695 |
| LINC02754 | 1.40632 | 4.16207 | 6.09099 | 3.18E-05 | 0.005088259 | 2.55713 |
| SV2B | 1.86720 | 1.61445 | 6.06109 | 3.21E-05 | 0.005088259 | 2.62651 |
| SATL1 | 2.03346 | 1.97507 | 6.05453 | 3.25E-05 | 0.005088259 | 2.52319 |
| GUSBP3 | 1.98799 | 2.71806 | 6.05236 | 3.26E-05 | 0.005088259 | 2.64960 |
| LINC01427 | 1.92934 | 3.99278 | 6.07421 | 3.27E-05 | 0.005088259 | 2.57210 |
| KRT78 | 2.25643 | 2.11325 | 6.05866 | 3.35E-05 | 0.005139948 | 2.62889 |
| PPP3CB-AS1 | 1.59230 | 2.96754 | 6.03053 | 3.38E-05 | 0.005139948 | 2.60698 |
| AQP7 | 1.68813 | 3.30853 | 6.02088 | 3.44E-05 | 0.005139948 | 2.60119 |
| ZBED3-AS1 | 1.60850 | 4.41013 | 6.04118 | 3.45E-05 | 0.005139948 | 2.45010 |
| MROH7 | 2.28896 | 1.69513 | 6.01821 | 3.45E-05 | 0.005139948 | 2.53363 |
| NR2F2-AS1 | 1.83392 | 2.09536 | 6.00427 | 3.54E-05 | 0.005162888 | 2.55804 |
| LINC01226 | 1.59577 | 2.51834 | 6.00096 | 3.56E-05 | 0.005162888 | 2.56799 |
| SPP2 | 1.46804 | 2.65075 | 5.99995 | 3.56E-05 | 0.005162888 | 2.56646 |
| TERB1 | 1.87574 | 2.88422 | 6.01244 | 3.62E-05 | 0.005205423 | 2.54834 |
| L3MBTL1 | 1.47488 | 4.13398 | 5.99755 | 3.71E-05 | 0.005215988 | 2.41093 |
| ARRDC3-AS1 | 1.38559 | 2.95048 | 5.97474 | 3.72E-05 | 0.005215988 | 2.51950 |
| FAM153CP | 1.74676 | 3.43771 | 5.99348 | 3.74E-05 | 0.005215988 | 2.47848 |
| ASB14 | 2.10133 | 2.25537 | 5.96881 | 3.75E-05 | 0.005215988 | 2.49716 |
| LINC01562 | 2.08527 | 1.85825 | 5.96081 | 3.81E-05 | 0.005223668 | 2.47876 |
| NIBAN3 | 1.56672 | 2.66981 | 5.95821 | 3.82E-05 | 0.005223668 | 2.49571 |
| TNPO1P3 | 3.74137 | 0.31189 | 5.94389 | 3.92E-05 | 0.005308856 | 1.70533 |
| RBMS3-AS3 | 1.90104 | 2.01343 | 5.93612 | 3.97E-05 | 0.005336048 | 2.45012 |
| NEGR1 | 1.69667 | 4.03089 | 5.95041 | 4.02E-05 | 0.005342851 | 2.36675 |
| LINC01515 | 1.53280 | 3.07733 | 5.92152 | 4.07E-05 | 0.005342851 | 2.42925 |
| LHFPL5 | 2.11782 | 1.59613 | 5.92134 | 4.07E-05 | 0.005342851 | 2.39918 |
| ITFG2 | 1.24528 | 3.69907 | 5.90841 | 4.16E-05 | 0.005419233 | 2.36498 |
| MYT1L | 1.82941 | 2.64331 | 5.89583 | 4.25E-05 | 0.005493897 | 2.39957 |
| GSTO2 | 1.82147 | 3.24897 | 5.88466 | 4.33E-05 | 0.005513219 | 2.38143 |
| CCDC18-AS1 | 1.52918 | 4.34779 | 5.89842 | 4.39E-05 | 0.005513219 | 2.21951 |
| SMIM11A | 1.91885 | 1.84480 | 5.87628 | 4.40E-05 | 0.005513219 | 2.32314 |
| RAD51B | 1.44805 | 3.32435 | 5.87517 | 4.40E-05 | 0.005513219 | 2.34536 |
| SCART1 | 1.93609 | 3.49276 | 5.89298 | 4.43E-05 | 0.005513219 | 2.32079 |
| SOX9-AS1 | 1.83660 | 2.12791 | 5.86543 | 4.48E-05 | 0.005518514 | 2.34923 |
| SCYL2P1 | 3.24652 | -0.46288 | 5.86114 | 4.51E-05 | 0.005518514 | 1.59417 |
| PRSS27 | 2.68390 | 1.70104 | 5.87529 | 4.57E-05 | 0.005518514 | 2.28140 |
| HOPX | 2.26033 | 3.03220 | 5.87466 | 4.57E-05 | 0.005518514 | 2.32775 |
| LINC01356 | 1.81394 | 1.16905 | 5.84433 | 4.64E-05 | 0.005518514 | 2.17399 |
| UBAP1L | 1.93328 | 2.37725 | 5.84378 | 4.65E-05 | 0.005518514 | 2.30899 |
| CCDC144A | 2.01416 | 3.31412 | 5.86244 | 4.67E-05 | 0.005518514 | 2.29177 |
| SPIN3 | 1.43776 | 4.38743 | 5.85277 | 4.74E-05 | 0.005570603 | 2.14970 |
| IFNE | 2.23221 | 1.54332 | 5.82556 | 4.87E-05 | 0.005676208 | 2.25637 |
| SULT1A3 | 2.67240 | 3.01539 | 5.80318 | 5.00E-05 | 0.005740369 | 2.12979 |
| SND1-IT1 | 2.44331 | 0.63606 | 5.79616 | 5.04E-05 | 0.005740369 | 2.04588 |
| C5orf63 | 1.55642 | 2.82228 | 5.78246 | 5.16E-05 | 0.005740369 | 2.21142 |
| MAGI1-IT1 | 2.12777 | 1.20557 | 5.78154 | 5.17E-05 | 0.005740369 | 2.16845 |
| C12orf50 | 1.54124 | 2.53032 | 5.77988 | 5.19E-05 | 0.005740369 | 2.20803 |
| LINC00923 | 1.63125 | 3.49178 | 5.79919 | 5.20E-05 | 0.005740369 | 2.14144 |
| CSTF3-DT | 2.36586 | 0.82853 | 5.77704 | 5.21E-05 | 0.005740369 | 2.03169 |
| ZNF846 | 1.43960 | 4.29886 | 5.79718 | 5.22E-05 | 0.005740369 | 2.06614 |
| SGCE | 1.63878 | 1.82683 | 5.77463 | 5.23E-05 | 0.005740369 | 2.19987 |
| ABCA11P | 1.49445 | 2.20399 | 5.76228 | 5.35E-05 | 0.005812003 | 2.18006 |
| FAM228B | 1.28027 | 4.35572 | 5.78041 | 5.37E-05 | 0.005812003 | 2.01630 |
| FRRS1L | 1.96661 | 2.56771 | 5.77325 | 5.44E-05 | 0.005845713 | 2.16841 |
| MRPL45P2 | 1.94167 | 1.81993 | 5.74533 | 5.51E-05 | 0.005859542 | 2.13194 |
| ZNF571-AS1 | 1.44219 | 2.70485 | 5.74398 | 5.52E-05 | 0.005859542 | 2.14887 |
| ZNF790-AS1 | 1.31414 | 3.13546 | 5.72927 | 5.66E-05 | 0.005972503 | 2.08624 |
| GUSBP2 | 1.40796 | 3.33036 | 5.71797 | 5.77E-05 | 0.006052233 | 2.07061 |
| SERPINI2 | 1.55150 | 3.67574 | 5.72451 | 5.91E-05 | 0.006157376 | 2.01437 |
| ABCA9 | 1.64313 | 2.06514 | 5.69805 | 5.98E-05 | 0.006186967 | 2.07341 |
| DANT2 | 1.94076 | 2.58509 | 5.70222 | 6.07E-05 | 0.006243386 | 2.06377 |
| PARP15 | 1.68721 | 3.74097 | 5.70566 | 6.10E-05 | 0.006243386 | 1.99333 |
| LINC01285 | 1.86801 | 1.61212 | 5.67943 | 6.17E-05 | 0.006248086 | 2.00015 |
| RAD51D | 1.17023 | 3.49054 | 5.67826 | 6.18E-05 | 0.006248086 | 1.97021 |
| RABGAP1L-DT | 1.57962 | 2.82725 | 5.66902 | 6.28E-05 | 0.006272095 | 2.01176 |
| ANKRD20A11P | 1.93461 | 2.65853 | 5.66842 | 6.29E-05 | 0.006272095 | 2.02666 |
| L3MBTL4-AS1 | 2.16814 | 1.47281 | 5.66252 | 6.36E-05 | 0.006272095 | 1.91505 |
| TAS2R4 | 1.85588 | 2.29510 | 5.66227 | 6.36E-05 | 0.006272095 | 2.01367 |
| MIR3936HG | 1.51907 | 2.94654 | 5.64189 | 6.59E-05 | 0.006440347 | 1.96158 |
| PDE6A | 1.91208 | 3.34257 | 5.65990 | 6.61E-05 | 0.006440347 | 1.97645 |
| RNF217-AS1 | 1.77565 | 1.92057 | 5.62866 | 6.74E-05 | 0.006534341 | 1.91979 |
| LRRC37A3 | 1.40585 | 3.97479 | 5.63322 | 6.91E-05 | 0.006655149 | 1.84053 |
| ARHGEF26-AS1 | 1.36256 | 3.59743 | 5.60727 | 7.00E-05 | 0.006705057 | 1.85929 |
| LINC00343 | 1.61896 | 3.27718 | 5.61354 | 7.13E-05 | 0.006789273 | 1.87229 |
| CBWD6 | 1.71753 | 3.20704 | 5.59163 | 7.22E-05 | 0.006841162 | 1.88886 |
| TNS1-AS1 | 1.90475 | 1.63441 | 5.58481 | 7.28E-05 | 0.006855454 | 1.88439 |
| LINC02027 | 1.85159 | 2.50314 | 5.58148 | 7.32E-05 | 0.0068568 | 1.86244 |
| FRG1-DT | 1.58518 | 2.79278 | 5.57304 | 7.43E-05 | 0.006901198 | 1.85700 |
| LINC00456 | 2.11982 | 0.89920 | 5.57142 | 7.45E-05 | 0.006901198 | 1.67988 |
| CDKL3 | 1.56442 | 1.85359 | 5.55688 | 7.64E-05 | 0.006998654 | 1.83850 |
| FRA10AC1 | 1.43701 | 3.36536 | 5.55338 | 7.69E-05 | 0.006998654 | 1.79936 |
| GLB1L3 | 1.95997 | 0.56942 | 5.55332 | 7.69E-05 | 0.006998654 | 1.68534 |
| LINC00641 | 1.52537 | 3.40110 | 5.56722 | 7.76E-05 | 0.006998654 | 1.76530 |
| KLRD1 | 1.58931 | 3.43802 | 5.56298 | 7.82E-05 | 0.006998654 | 1.75124 |
| LINC00997 | 1.43801 | 1.97193 | 5.53851 | 7.89E-05 | 0.006998654 | 1.81455 |
| EID3 | 2.00868 | 1.09990 | 5.53321 | 7.97E-05 | 0.006998654 | 1.69954 |
| MSANTD2 | 1.50592 | 2.73604 | 5.52815 | 8.04E-05 | 0.006998654 | 1.78972 |
| LNC-LBCS | 0.89349 | 4.42588 | 5.52781 | 8.04E-05 | 0.006998654 | 1.63219 |
| LINC01579 | 2.14603 | 1.42098 | 5.52774 | 8.05E-05 | 0.006998654 | 1.73464 |
| RYR3 | 1.94983 | 1.02075 | 5.52651 | 8.06E-05 | 0.006998654 | 1.74245 |
| USP32P2 | 3.79211 | 0.24832 | 5.54462 | 8.07E-05 | 0.006998654 | 1.69212 |
| HLA-L | 1.75274 | 2.27856 | 5.52163 | 8.13E-05 | 0.006998654 | 1.76159 |
| SMIM5 | 1.03168 | 4.49767 | 5.52077 | 8.14E-05 | 0.006998654 | 1.61033 |
| PDLIM3 | 3.05170 | -0.09577 | 5.51542 | 8.22E-05 | 0.007028599 | 1.28643 |
| PJVK | 1.79068 | 1.46447 | 5.50646 | 8.35E-05 | 0.007103996 | 1.74758 |
| LSP1P4 | 2.08375 | 1.17023 | 5.49539 | 8.52E-05 | 0.007170729 | 1.61117 |
| GHRLOS | 1.80751 | 1.41829 | 5.49411 | 8.54E-05 | 0.007170729 | 1.71631 |
| PSEN2 | 1.56269 | 4.42376 | 5.50888 | 8.59E-05 | 0.007170729 | 1.56790 |
| AMY2B | 1.24534 | 3.07997 | 5.48798 | 8.63E-05 | 0.007170729 | 1.69107 |
| SRP14-AS1 | 1.49530 | 2.32172 | 5.47895 | 8.77E-05 | 0.007170729 | 1.71582 |
| DNAJC19P5 | 2.11972 | -0.05187 | 5.47594 | 8.81E-05 | 0.007170729 | 1.55449 |
| LAIR1 | 1.78631 | 2.69017 | 5.49297 | 8.83E-05 | 0.007170729 | 1.69612 |
| PPP5D1 | 2.03091 | 0.72865 | 5.46809 | 8.94E-05 | 0.007170729 | 1.60350 |
| TIMM23B | 1.58270 | 4.07082 | 5.48623 | 8.94E-05 | 0.007170729 | 1.57966 |
| RALY-AS1 | 1.42625 | 3.47345 | 5.46972 | 8.98E-05 | 0.007170729 | 1.62426 |
| ODAPH | 2.38412 | 1.45718 | 5.46522 | 8.98E-05 | 0.007170729 | 1.49450 |
| OR14J1 | 1.69649 | 4.39773 | 5.48273 | 8.99E-05 | 0.007170729 | 1.47266 |
| MIR3648-1 | -1.30087 | 5.37965 | -5.47950 | 9.04E-05 | 0.007170729 | 1.38240 |
| GJB1 | 1.75852 | 3.27674 | 5.47813 | 9.07E-05 | 0.007170729 | 1.66175 |
| IRS3P | 1.90816 | 1.98474 | 5.45937 | 9.08E-05 | 0.007170729 | 1.68301 |
| HDAC1P2 | 2.47497 | 0.13302 | 5.44980 | 9.23E-05 | 0.007216395 | 1.54598 |
| TCP10L | 1.81805 | 1.48906 | 5.44819 | 9.26E-05 | 0.007216395 | 1.62545 |
| TAS2R30 | 2.65323 | 0.80514 | 5.44779 | 9.26E-05 | 0.007216395 | 1.50501 |
| LINC02254 | 1.38954 | 4.11083 | 5.45459 | 9.45E-05 | 0.007296139 | 1.47019 |
| C21orf62-AS1 | 1.51307 | 2.73457 | 5.43633 | 9.45E-05 | 0.007296139 | 1.62778 |
| CLHC1 | 1.27123 | 4.77461 | 5.44381 | 9.63E-05 | 0.007398053 | 1.41100 |
| CAPN8 | 1.36462 | 3.93266 | 5.44070 | 9.68E-05 | 0.007404541 | 1.48911 |
| ZDHHC15 | 2.54143 | 0.21418 | 5.41698 | 9.78E-05 | 0.007447181 | 1.21527 |
| COX10-AS1 | 1.31250 | 3.82541 | 5.42323 | 9.84E-05 | 0.007456593 | 1.48700 |
| NKAPP1 | 1.54633 | 1.96249 | 5.40911 | 9.92E-05 | 0.007483574 | 1.59825 |
| DPY19L2P1 | 1.64235 | 1.35784 | 5.40344 | 0.000100209 | 0.007507233 | 1.48698 |
| CFAP299 | 1.30097 | 3.99791 | 5.41327 | 0.000100421 | 0.007507233 | 1.45701 |
| ZNF213-AS1 | 1.44425 | 4.02301 | 5.41380 | 0.000101524 | 0.007529032 | 1.40761 |
| LINC02615 | 1.91690 | 2.70555 | 5.40978 | 0.000102246 | 0.007529032 | 1.56947 |
| DET1 | 1.12451 | 3.22408 | 5.39149 | 0.00010236 | 0.007529032 | 1.53019 |
| RPL32P3 | 1.25914 | 4.51714 | 5.40659 | 0.000102823 | 0.007529032 | 1.34169 |
| ZNF169 | 1.69607 | 2.50193 | 5.38814 | 0.00010297 | 0.007529032 | 1.55438 |
| SH3BP5-AS1 | 1.46403 | 3.51158 | 5.38580 | 0.000106666 | 0.007732951 | 1.44444 |
| LYPLAL1-AS1 | 1.91166 | 1.66639 | 5.36821 | 0.000106687 | 0.007732951 | 1.51934 |
| LINC00547 | 1.76321 | 1.49989 | 5.36376 | 0.000107536 | 0.007760732 | 1.51316 |
| LINC01301 | 1.68498 | 1.42844 | 5.35828 | 0.00010859 | 0.007781759 | 1.50510 |
| OLA1P2 | 3.00395 | 0.19358 | 5.35657 | 0.000108921 | 0.007781759 | 1.11161 |
| IL33 | -1.20691 | 5.86681 | -5.36986 | 0.000109717 | 0.007781759 | 1.25836 |
| C2orf92 | 1.49552 | 3.04496 | 5.35966 | 0.00010976 | 0.007781759 | 1.45011 |
| HMGA2 | 1.13330 | 5.05326 | 5.36757 | 0.000110161 | 0.007781759 | 1.25897 |
| GUSBP1 | 1.40908 | 3.09400 | 5.34542 | 0.000111108 | 0.007792013 | 1.44013 |
| LINC02013 | 3.13633 | 0.16381 | 5.34475 | 0.000111241 | 0.007792013 | 1.03772 |
| LINC00355 | 2.13989 | 0.33716 | 5.33551 | 0.000113089 | 0.007856319 | 1.37954 |
| EEF1A1P3 | 2.45512 | 0.57562 | 5.33545 | 0.000113102 | 0.007856319 | 1.23649 |
| PVT1 | 1.64329 | 4.42919 | 5.34770 | 0.000114106 | 0.007878265 | 1.28199 |
| LINC01695 | 1.81492 | 1.78072 | 5.32924 | 0.000114363 | 0.007878265 | 1.46507 |
| MAMDC2-AS1 | 2.45707 | 0.91764 | 5.32643 | 0.000114938 | 0.007885293 | 1.14942 |
| DDR1-DT | 1.75026 | 2.31150 | 5.32345 | 0.000115551 | 0.007894863 | 1.45526 |
| CASC19 | 1.96763 | 3.72112 | 5.33462 | 0.000116783 | 0.007946467 | 1.35767 |
| LINC00939 | 1.48821 | 3.21369 | 5.31899 | 0.000117996 | 0.007996391 | 1.38109 |
| THUMPD3-AS1 | 1.22329 | 4.39956 | 5.32394 | 0.000119018 | 0.00801264 | 1.22413 |
| LINC01359 | 1.98811 | 0.93501 | 5.30599 | 0.000119214 | 0.00801264 | 1.35586 |
| LAMTOR5-AS1 | 1.68665 | 2.54015 | 5.30382 | 0.000119678 | 0.00801264 | 1.41872 |
| OR1A1 | 1.98262 | 2.42499 | 5.30368 | 0.00012017 | 0.008013444 | 1.41963 |
| ALOX12P2 | 1.40449 | 4.41553 | 5.30316 | 0.000123494 | 0.008171491 | 1.15506 |
| UBE3D | 1.25940 | 3.53914 | 5.28616 | 0.000123521 | 0.008171491 | 1.32071 |
| USHBP1 | 1.67685 | 1.55994 | 5.27782 | 0.00012538 | 0.00826172 | 1.37210 |
| ACADL | 1.74113 | 2.69868 | 5.28354 | 0.000126388 | 0.008295357 | 1.35848 |
| LIMD1-AS1 | 2.32941 | 1.67649 | 5.26522 | 0.000128245 | 0.00838419 | 1.24152 |
| ZNF789 | 1.57647 | 2.52305 | 5.26144 | 0.000129117 | 0.008408268 | 1.34334 |
| ZFHX2-AS1 | 1.88815 | 1.69116 | 5.25782 | 0.000129959 | 0.008430146 | 1.32682 |
| PTOV1-AS1 | 1.73177 | 1.28609 | 5.25166 | 0.000131406 | 0.00847144 | 1.28549 |
| EDIL3-DT | 1.76260 | 2.26463 | 5.25079 | 0.000131612 | 0.00847144 | 1.33418 |
| KCNV1 | 2.27710 | 1.43603 | 5.22912 | 0.000136839 | 0.008774038 | 1.16306 |
| NXNL2 | 2.59009 | 0.74734 | 5.22497 | 0.000137864 | 0.008785575 | 1.12138 |
| FLVCR1-DT | 1.86108 | 1.81647 | 5.22413 | 0.000138073 | 0.008785575 | 1.26694 |
| DRAIC | 1.40045 | 3.73279 | 5.22814 | 0.000139258 | 0.00881414 | 1.19410 |
| RN7SL832P | 2.80093 | -0.80676 | 5.21810 | 0.00013958 | 0.00881414 | 0.74306 |
| AFF3 | 2.39655 | 0.92216 | 5.20544 | 0.000142798 | 0.008983339 | 1.05027 |
| LINC02887 | 2.34158 | 0.68741 | 5.20884 | 0.000143709 | 0.00899732 | 1.22570 |
| STRIP2 | 1.49628 | 2.73204 | 5.19721 | 0.000144932 | 0.00899732 | 1.22056 |
| GIMAP2 | 1.93081 | 2.11857 | 5.20722 | 0.000145991 | 0.00899732 | 1.23848 |
| LINC01967 | 1.51173 | 1.88034 | 5.19245 | 0.000146182 | 0.00899732 | 1.23621 |
| LINC01843 | 1.57768 | 1.17454 | 5.19171 | 0.000146378 | 0.00899732 | 1.20466 |
| CAPN3 | 1.49263 | 3.54143 | 5.20776 | 0.000146412 | 0.00899732 | 1.12179 |
| CSAD | 1.56131 | 3.55327 | 5.20629 | 0.000146798 | 0.00899732 | 1.11988 |
| POLR2J4 | 1.30625 | 3.37748 | 5.18628 | 0.000147818 | 0.009026638 | 1.14633 |
| WFDC1 | 1.99325 | 1.78407 | 5.17819 | 0.000149994 | 0.009086668 | 1.21044 |
| ADAMTSL4-AS1 | 1.84227 | 1.05749 | 5.17719 | 0.000150263 | 0.009086668 | 1.17614 |
| LINC01764 | 1.99293 | 1.04739 | 5.17656 | 0.000150436 | 0.009086668 | 1.15791 |
| OR2T12 | 1.60224 | 2.16830 | 5.16683 | 0.000153103 | 0.009214374 | 1.18980 |
| EYS | 2.27155 | 1.72057 | 5.15836 | 0.000155465 | 0.009322838 | 1.13808 |
| LINC02042 | 1.92243 | 0.26285 | 5.15414 | 0.000156657 | 0.009360689 | 1.00401 |
| SCML4 | 1.56385 | 2.16283 | 5.13946 | 0.000160873 | 0.009534134 | 1.14578 |
| LINC01361 | 2.65868 | -0.13392 | 5.13780 | 0.000161358 | 0.009534134 | 0.67690 |
| LINC02840 | 1.31917 | 2.12142 | 5.13598 | 0.000161891 | 0.009534134 | 1.13385 |
| SCN9A | 1.30579 | 2.91308 | 5.13416 | 0.000162424 | 0.009534134 | 1.07107 |
| CYP1A2 | 1.82144 | 1.32017 | 5.13336 | 0.000162662 | 0.009534134 | 1.06077 |
| MSH5 | 1.27617 | 4.19674 | 5.14800 | 0.000162991 | 0.009534134 | 0.92607 |
| MYLK-AS1 | 1.90034 | 1.23006 | 5.12581 | 0.0001649 | 0.009595831 | 1.09307 |
| ABCA6 | 1.35230 | 2.14855 | 5.12482 | 0.000165197 | 0.009595831 | 1.12200 |
| MAFTRR | 3.15580 | -0.53188 | 5.11954 | 0.000166787 | 0.009646681 | 0.56612 |
| SMC5-AS1 | 1.53677 | 1.89243 | 5.11808 | 0.00016723 | 0.009646681 | 1.11093 |
| OR2V2 | 2.45059 | 0.83912 | 5.11426 | 0.000168393 | 0.00967695 | 0.92272 |
| SRARP | 1.86421 | 1.14336 | 5.11255 | 0.000168916 | 0.00967695 | 1.07057 |
| RBFADN | 1.81711 | 3.07971 | 5.12355 | 0.000170328 | 0.009724424 | 1.03641 |
| THEM4 | 0.87213 | 3.82292 | 5.10023 | 0.000172735 | 0.009793907 | 0.91483 |
| LINC01239 | 1.48537 | 1.66950 | 5.09532 | 0.000174283 | 0.009793907 | 1.07204 |
| PSMG4 | 1.42323 | 2.95744 | 5.09526 | 0.0001743 | 0.009793907 | 1.01997 |
| CROCCP3 | 1.26306 | 4.38703 | 5.11070 | 0.000174317 | 0.009793907 | 0.82808 |
| LINC00643 | 1.12380 | 2.95173 | 5.09297 | 0.000175029 | 0.009793907 | 1.01095 |
| LINC02268 | 1.68303 | 2.40925 | 5.09064 | 0.00017577 | 0.009793907 | 1.06256 |
| ZKSCAN2-DT | 2.09988 | 1.07751 | 5.08948 | 0.000176141 | 0.009793907 | 1.03835 |
| EIF1B-AS1 | 1.28016 | 4.14973 | 5.10461 | 0.000176244 | 0.009793907 | 0.85318 |
| FAM153A | 2.13807 | 1.67875 | 5.08644 | 0.000177116 | 0.009809665 | 0.99530 |
| TMC2 | 3.52125 | 0.16553 | 5.08136 | 0.00017876 | 0.009867897 | 0.36187 |
| LINC02595 | 2.38031 | 1.37142 | 5.08635 | 0.000179815 | 0.009893398 | 1.01555 |
| LINC00894 | 1.76995 | 3.98807 | 5.08655 | 0.000182088 | 0.009985508 | 0.98612 |
| PPIEL | 1.72283 | 2.40303 | 5.06872 | 0.000182921 | 0.009998266 | 1.01158 |
| ADCY4 | 1.72535 | 0.40739 | 5.06526 | 0.000184075 | 0.010023907 | 0.94807 |
| USP2-AS1 | 1.37694 | 3.83373 | 5.07830 | 0.000184823 | 0.010023907 | 0.87655 |
| CYP4A22-AS1 | 1.34884 | 2.51963 | 5.06039 | 0.000185716 | 0.010023907 | 0.99892 |
| TAS2R14 | 2.11273 | 1.04254 | 5.06015 | 0.000185795 | 0.010023907 | 0.98745 |
| SCAND2P | 1.34468 | 3.95536 | 5.07193 | 0.000186964 | 0.010051944 | 0.80981 |
| ADGRF2 | 1.64169 | 2.51080 | 5.05508 | 0.000187521 | 0.010051944 | 0.99501 |
| TSEN15 | 0.85829 | 4.25362 | 5.05054 | 0.000189077 | 0.010076388 | 0.80727 |
| GPR52 | 2.55128 | -0.20763 | 5.04995 | 0.000189282 | 0.010076388 | 0.71054 |
| DLGAP1-AS2 | 1.57410 | 2.30535 | 5.04699 | 0.000190305 | 0.010076388 | 0.98815 |
| LINC00652 | 1.55588 | 1.85002 | 5.04673 | 0.000190394 | 0.010076388 | 0.98543 |
| SEM1 | 1.14132 | 3.51003 | 5.04315 | 0.000191643 | 0.010104135 | 0.89642 |
| BMS1P2 | 2.15546 | 2.66812 | 5.05687 | 0.000192131 | 0.010104135 | 0.98475 |
| MUC5AC | -4.19710 | 9.84584 | -5.05246 | 0.00019367 | 0.010104844 | 0.93927 |
| B4GALNT2 | 1.35221 | 4.29836 | 5.05219 | 0.000193767 | 0.010104844 | 0.72988 |
| SRSF12 | 2.17993 | 1.48058 | 5.03618 | 0.000194093 | 0.010104844 | 0.94179 |
| SLC9C2 | 1.69901 | 2.68381 | 5.03484 | 0.000194569 | 0.010104844 | 0.96555 |
| ARMCX4 | 1.41693 | 3.10293 | 5.02950 | 0.000198431 | 0.010273445 | 0.87584 |
| CNN1 | 2.03343 | 0.34133 | 5.02060 | 0.000199692 | 0.0102976 | 0.77036 |
| CYP46A1 | 1.57484 | 1.65044 | 5.01788 | 0.000200688 | 0.0102976 | 0.94010 |
| DCN | 3.78265 | -0.80889 | 5.01740 | 0.000200862 | 0.0102976 | 0.13722 |
| MIR100HG | 1.55671 | 3.11667 | 5.01602 | 0.000201369 | 0.0102976 | 0.90671 |
| CLEC3A | 2.46155 | -0.64573 | 5.00928 | 0.000203867 | 0.010375832 | 0.51879 |
| PACRGL | 0.88448 | 4.65530 | 5.00825 | 0.000204248 | 0.010375832 | 0.65604 |
| LGALSL-DT | 2.57619 | 1.11338 | 5.00707 | 0.000204766 | 0.010375832 | 0.83591 |
| STPG2-AS1 | 2.00785 | 1.20950 | 4.99790 | 0.000208151 | 0.010497759 | 0.79715 |
| SLC4A1 | 2.23456 | 0.67836 | 4.99616 | 0.000208814 | 0.010497759 | 0.81348 |
| FOSB | 2.09548 | 2.52509 | 5.01032 | 0.000209061 | 0.010497759 | 0.90101 |
| RNASEH2B-AS1 | 1.68103 | 2.72987 | 5.00636 | 0.00021057 | 0.010537863 | 0.85852 |
| RPARP-AS1 | 1.30160 | 3.96686 | 5.00435 | 0.000211341 | 0.010537863 | 0.72659 |
| DNAJC17 | 1.13658 | 3.01132 | 4.98852 | 0.000211756 | 0.010537863 | 0.83759 |
| LINC00670 | 1.91410 | 1.57034 | 4.98636 | 0.000212594 | 0.010548064 | 0.86739 |
| RNF213-AS1 | 1.08820 | 4.23887 | 4.99885 | 0.000213466 | 0.010559895 | 0.62527 |
| LINC01907 | 2.10195 | 1.65786 | 4.97911 | 0.000215436 | 0.01062582 | 0.86410 |
| LINC01588 | 1.29768 | 3.25266 | 4.97137 | 0.000218514 | 0.010719052 | 0.76214 |
| CRYBB2P1 | 1.58737 | 3.53312 | 4.98575 | 0.000218612 | 0.010719052 | 0.70584 |
| CYP2F2P | 1.51835 | 3.17671 | 4.96693 | 0.000220298 | 0.010768294 | 0.84771 |
| KANTR | 1.30294 | 3.19488 | 4.96542 | 0.000220908 | 0.010768294 | 0.80076 |
| EP400P1 | 1.18578 | 3.99805 | 4.96683 | 0.000226276 | 0.010997792 | 0.62809 |
| FAM95C | 1.41882 | 3.10355 | 4.94778 | 0.000228177 | 0.011004315 | 0.76677 |
| MTFR2P1 | 2.02080 | 0.16916 | 4.94680 | 0.000228587 | 0.011004315 | 0.61970 |
| KBTBD12 | 1.80764 | 2.99741 | 4.96062 | 0.000228855 | 0.011004315 | 0.77933 |
| LINC02604 | 1.97028 | 2.56135 | 4.94937 | 0.00022905 | 0.011004315 | 0.81934 |
| PLAT | 1.46887 | 4.33392 | 4.95792 | 0.000229983 | 0.011017384 | 0.60188 |
| MCM3AP-AS1 | 1.82395 | 1.81068 | 4.94018 | 0.00023138 | 0.01103582 | 0.80869 |
| LRRC37A | 1.22635 | 3.41637 | 4.93945 | 0.000231692 | 0.01103582 | 0.68699 |
| U2AF1L4 | 1.39862 | 2.64456 | 4.93732 | 0.000232599 | 0.011047473 | 0.77082 |
| MIAT | 2.08156 | 0.99317 | 4.93246 | 0.000234688 | 0.011097699 | 0.63759 |
| LINC01089 | 1.62982 | 2.62062 | 4.94422 | 0.000234988 | 0.011097699 | 0.75788 |
| TRDMT1 | 0.99216 | 4.32589 | 4.92392 | 0.000238398 | 0.011181259 | 0.56270 |
| ANKRD18A | 1.40387 | 3.69935 | 4.93640 | 0.000239196 | 0.011181259 | 0.64239 |
| TMCC1-AS1 | 1.25883 | 2.83175 | 4.92196 | 0.000239261 | 0.011181259 | 0.72843 |
| SPC25 | 1.95270 | 2.32289 | 4.93584 | 0.00023944 | 0.011181259 | 0.77082 |
| ERVK9-11 | 1.48897 | 1.93532 | 4.91786 | 0.000241073 | 0.011198684 | 0.76785 |
| RPL34-DT | 1.76309 | 0.26978 | 4.91767 | 0.000241157 | 0.011198684 | 0.68973 |
| ST7-OT4 | 1.89051 | 0.40117 | 4.91611 | 0.000241847 | 0.01119954 | 0.68024 |
| NCK1-DT | 1.29816 | 3.05019 | 4.91259 | 0.000243419 | 0.01123102 | 0.70947 |
| PPP1R26-AS1 | 1.31366 | 2.32193 | 4.91158 | 0.000243874 | 0.01123102 | 0.73971 |
| ANKRD40CL | 2.65388 | -0.09150 | 4.90589 | 0.000246443 | 0.011318037 | 0.45989 |
| SNHG27 | 2.58335 | 0.42993 | 4.90237 | 0.000248044 | 0.011360259 | 0.60933 |
| FBXO30-DT | 0.96163 | 4.32062 | 4.89701 | 0.000250504 | 0.011441493 | 0.46063 |
| RBM5-AS1 | 2.44425 | -0.51741 | 4.89287 | 0.000252424 | 0.011497705 | 0.44560 |
| SLC16A1-AS1 | 1.60351 | 2.50511 | 4.89834 | 0.00025338 | 0.011509793 | 0.69116 |
| RBM22P2 | 1.74718 | 2.13165 | 4.89220 | 0.00025433 | 0.011521582 | 0.71333 |
| ZDHHC11 | 1.38279 | 3.08834 | 4.89307 | 0.000256324 | 0.011550994 | 0.61365 |
| ANKRD36BP2 | 1.30856 | 3.62713 | 4.89850 | 0.000256365 | 0.011550994 | 0.53980 |
| C9orf147 | 2.23825 | 1.12401 | 4.88194 | 0.00025756 | 0.011573538 | 0.61996 |
| SIGLEC10 | 1.82147 | 1.91029 | 4.88274 | 0.000258875 | 0.011601364 | 0.70212 |
| SNHG17 | 1.22435 | 3.79526 | 4.89104 | 0.00025989 | 0.011615619 | 0.49881 |
| MTCYBP28 | 3.06581 | -1.35015 | 4.87160 | 0.000262521 | 0.01170183 | -0.05386 |
| PIGL | 1.12186 | 4.67651 | 4.88157 | 0.000264441 | 0.011725969 | 0.35170 |
| LINC01341 | 1.85439 | 1.04657 | 4.86759 | 0.000264469 | 0.011725969 | 0.67385 |
| LINC02275 | 2.34598 | 0.19259 | 4.86525 | 0.000265613 | 0.011745452 | 0.28342 |
| ATP5F1BP1 | 2.85301 | -0.59630 | 4.85870 | 0.000268845 | 0.011827889 | 0.22541 |
| DLEU2 | 1.56356 | 3.62233 | 4.87246 | 0.000268896 | 0.011827889 | 0.50812 |
| TTLL7-IT1 | 1.87949 | 1.47479 | 4.85347 | 0.000271455 | 0.011909005 | 0.63586 |
| GVINP1 | 2.00904 | 1.43729 | 4.85110 | 0.000272648 | 0.011929957 | 0.62722 |
| ZNF226 | 1.03654 | 4.56221 | 4.85602 | 0.000273889 | 0.011952883 | 0.38822 |
| RUNDC3A | 1.91942 | 1.10080 | 4.84487 | 0.000275806 | 0.011977475 | 0.56541 |
| SDHAP1 | 1.20563 | 3.79598 | 4.85848 | 0.000275889 | 0.011977475 | 0.40421 |
| OR52A1 | 1.49587 | 1.98301 | 4.84308 | 0.000276721 | 0.011982396 | 0.62625 |
| NPIPB2 | 1.60283 | 1.30233 | 4.83560 | 0.000280574 | 0.012117755 | 0.62209 |
| ANKRD39 | 1.23683 | 3.45254 | 4.82977 | 0.000283619 | 0.012217587 | 0.53347 |
| CLSPN | 1.59019 | 3.12369 | 4.84126 | 0.000284757 | 0.012234995 | 0.54457 |
| GNRHR2 | 1.70591 | 2.73337 | 4.82369 | 0.000286827 | 0.012289002 | 0.61019 |
| PI4KAP1 | 1.31836 | 2.97145 | 4.83493 | 0.000288094 | 0.012289002 | 0.45245 |
| NAIP | 1.45767 | 4.62998 | 4.83398 | 0.000288594 | 0.012289002 | 0.33593 |
| CA13 | 0.82388 | 4.59580 | 4.81968 | 0.000288965 | 0.012289002 | 0.35297 |
| LINC01445 | 1.40854 | 2.05123 | 4.81831 | 0.000289699 | 0.012289002 | 0.58980 |
| LINC01446 | 2.64380 | -0.10649 | 4.80863 | 0.000294942 | 0.012407587 | 0.20406 |
| FGF13 | 1.49739 | 3.53687 | 4.82206 | 0.000294997 | 0.012407587 | 0.43967 |
| C8orf44 | 1.68770 | 2.16571 | 4.81001 | 0.000295107 | 0.012407587 | 0.57038 |
| ATP6AP1L | 1.52099 | 2.59007 | 4.80766 | 0.000295472 | 0.012407587 | 0.56438 |
| LINC02503 | 1.26845 | 4.98198 | 4.81595 | 0.000298332 | 0.01248333 | 0.17928 |
| MIR646HG | 1.63932 | 2.28315 | 4.79826 | 0.000300665 | 0.01248333 | 0.55015 |
| BVES | 2.33980 | 0.32549 | 4.79824 | 0.000300677 | 0.01248333 | 0.43648 |
| NLRC5 | -1.10280 | 3.91800 | -4.79769 | 0.000300985 | 0.01248333 | 0.39095 |
| PCDHGA12 | 1.73067 | 1.69142 | 4.79763 | 0.00030102 | 0.01248333 | 0.56560 |
| KCNB1 | 1.12236 | 3.59815 | 4.79334 | 0.000303425 | 0.012497427 | 0.37488 |
| GMDS-DT | 1.30355 | 4.78427 | 4.80452 | 0.000304682 | 0.012497427 | 0.23113 |
| SMG1P7 | 1.14048 | 2.51079 | 4.79110 | 0.000304683 | 0.012497427 | 0.51865 |
| TSIX | 1.33573 | 3.87111 | 4.80395 | 0.000305004 | 0.012497427 | 0.35724 |
| RDH16 | 1.71729 | 1.47347 | 4.79035 | 0.000305108 | 0.012497427 | 0.53570 |
| OR10H1 | 1.56509 | 2.06705 | 4.78122 | 0.000310325 | 0.012679989 | 0.53099 |
| LINC00884 | 1.25233 | 4.10790 | 4.79108 | 0.000312331 | 0.012730729 | 0.29622 |
| EBLN2 | 2.01041 | 1.38826 | 4.77414 | 0.000314431 | 0.012752676 | 0.51966 |
| DDX50P1 | 2.53010 | 0.86029 | 4.77388 | 0.000314585 | 0.012752676 | 0.28206 |
| UBL7-AS1 | 1.16409 | 3.63338 | 4.77855 | 0.000315164 | 0.012752676 | 0.34018 |
| GRAMD4P3 | 2.03349 | 2.09437 | 4.77994 | 0.000318411 | 0.012852843 | 0.51681 |
| SNX22 | 2.28006 | 0.54904 | 4.76146 | 0.000321927 | 0.012888242 | 0.37136 |
| VENTX | 2.42645 | 0.00272 | 4.75932 | 0.000323213 | 0.012888242 | 0.31721 |
| LACTB2-AS1 | 1.61886 | 3.05879 | 4.77213 | 0.000323449 | 0.012888242 | 0.42660 |
| CLEC12A-AS1 | 1.19465 | 3.38605 | 4.75875 | 0.000323554 | 0.012888242 | 0.35862 |
| LINC01443 | 2.16620 | 0.76677 | 4.75687 | 0.000324685 | 0.012888242 | 0.45373 |
| KLHL6-AS1 | 2.48579 | -0.20380 | 4.75635 | 0.000324999 | 0.012888242 | -0.03006 |
| HAND2-AS1 | 2.00646 | 1.53636 | 4.76180 | 0.000325097 | 0.012888242 | 0.49585 |
| ERICH1 | 1.16376 | 3.49387 | 4.75557 | 0.000325472 | 0.012888242 | 0.34588 |
| ALG13 | 1.01495 | 4.93240 | 4.76204 | 0.000329534 | 0.013018135 | 0.13590 |
| MEG3 | 2.43451 | 0.61831 | 4.74737 | 0.000330476 | 0.013024489 | 0.34012 |
| LINC01473 | 1.86448 | 1.14429 | 4.74592 | 0.00033137 | 0.01302893 | 0.44774 |
| CYP2B6 | 1.39739 | 2.15197 | 4.74395 | 0.000332586 | 0.013045982 | 0.44761 |
| MFF-DT | 1.71965 | 1.94147 | 4.73834 | 0.000336076 | 0.01314439 | 0.46338 |
| TPH2 | 1.74959 | 1.52457 | 4.73739 | 0.000336672 | 0.01314439 | 0.46181 |
| TPTE2P5 | 2.58931 | 0.45987 | 4.72826 | 0.000342448 | 0.013338665 | -0.04034 |
| UGT1A7 | 1.61960 | 0.96676 | 4.72392 | 0.000345229 | 0.013415658 | 0.43630 |
| AGAP1-IT1 | 2.30548 | -0.17185 | 4.72035 | 0.000347532 | 0.013473745 | 0.17837 |
| ZC3H12D | 1.62109 | 1.55357 | 4.71874 | 0.000348574 | 0.013482762 | 0.43030 |
| LINC-PINT | 1.59839 | 4.25206 | 4.72952 | 0.000349973 | 0.013500784 | 0.16358 |
| ZNF561-AS1 | 1.42220 | 3.15065 | 4.72780 | 0.000350659 | 0.013500784 | 0.32808 |
| CCDC162P | 1.09150 | 4.39911 | 4.72261 | 0.000354486 | 0.013591409 | 0.12137 |
| CTBP2P8 | 2.35114 | -0.03003 | 4.70949 | 0.000354644 | 0.013591409 | 0.23087 |
| PHYKPL | 1.21747 | 4.67068 | 4.72027 | 0.000356022 | 0.013612955 | 0.08168 |
| HNRNPA1P54 | 1.70535 | 3.30910 | 4.70577 | 0.000357458 | 0.013636567 | 0.39575 |
| OR5AS1 | 1.76313 | 2.93940 | 4.70482 | 0.000361405 | 0.013731207 | 0.38833 |
| SNHG26 | 1.43807 | 1.62897 | 4.69910 | 0.000361586 | 0.013731207 | 0.38761 |
| SPN | 1.57298 | 1.84730 | 4.69422 | 0.000364895 | 0.01382537 | 0.38548 |
| LIPC | 2.67158 | 0.10528 | 4.68972 | 0.000367973 | 0.013910377 | 0.17259 |
| ZNF586 | 1.28015 | 4.39696 | 4.69908 | 0.000370298 | 0.013966592 | 0.10728 |
| ZNF660 | 1.24929 | 2.16874 | 4.68438 | 0.000371661 | 0.013986366 | 0.33559 |
| LINC00106 | 2.65759 | -0.12042 | 4.68304 | 0.00037259 | 0.013989748 | 0.12508 |
| SRRM2-AS1 | 1.19763 | 2.29149 | 4.67784 | 0.000376233 | 0.014061025 | 0.32964 |
| TAS2R19 | 2.11242 | -0.20168 | 4.67736 | 0.000376567 | 0.014061025 | 0.15576 |
| RNASE7 | 2.40126 | 0.71721 | 4.67572 | 0.000377722 | 0.014061025 | 0.29632 |
| PNMA2 | 1.70371 | 1.44929 | 4.67553 | 0.000377862 | 0.014061025 | 0.32690 |
| INE1 | 2.11710 | 0.92903 | 4.67139 | 0.000380792 | 0.014138482 | 0.32208 |
| FOXL2NB | 1.34214 | 1.72298 | 4.66872 | 0.000382703 | 0.014148468 | 0.33846 |
| ZNF337-AS1 | 1.22467 | 3.03356 | 4.66864 | 0.000382758 | 0.014148468 | 0.23581 |
| CD226 | 1.48219 | 1.31078 | 4.66511 | 0.000385296 | 0.014210776 | 0.33515 |
| MITD1 | 1.05200 | 3.85065 | 4.66263 | 0.000387086 | 0.014245278 | 0.13043 |
| C12orf60 | 1.22934 | 2.06924 | 4.65316 | 0.00039401 | 0.01446816 | 0.31135 |
| LINC01234 | 2.58908 | 0.70790 | 4.64724 | 0.000398398 | 0.014575162 | 0.20895 |
| E2F3P2 | 1.37498 | 2.75790 | 4.64687 | 0.000398673 | 0.014575162 | 0.23838 |
| RN7SL67P | 3.15912 | -0.79154 | 4.64061 | 0.000405666 | 0.014776682 | 0.05001 |
| ZNF718 | 0.84735 | 3.40012 | 4.63721 | 0.000405958 | 0.014776682 | 0.13910 |
| ZNF382 | 1.09249 | 3.66788 | 4.62981 | 0.000411627 | 0.014950404 | 0.11491 |
| LINC02175 | 1.46562 | 1.55149 | 4.62816 | 0.000412902 | 0.014964097 | 0.27183 |
| PRICKLE4 | 2.84012 | -0.16268 | 4.63631 | 0.000416156 | 0.014978269 | 0.21505 |
| SULT1C2 | 2.10717 | 0.16841 | 4.62189 | 0.000417781 | 0.014978269 | 0.10080 |
| LINC01697 | 1.31160 | 2.82991 | 4.62155 | 0.00041805 | 0.014978269 | 0.18656 |
| BCO2 | 0.82786 | 4.94301 | 4.62128 | 0.000418262 | 0.014978269 | -0.09421 |
| RANBP17 | 1.20001 | 3.52452 | 4.62316 | 0.000418286 | 0.014978269 | 0.10776 |
| OR11A1 | 1.64097 | -0.00292 | 4.62074 | 0.000418684 | 0.014978269 | 0.23752 |
| AGAP9 | 1.65293 | 3.45080 | 4.63034 | 0.000420809 | 0.015005596 | 0.21371 |
| LDC1P | 1.90316 | -0.03881 | 4.61749 | 0.000421248 | 0.015005596 | 0.16080 |
| LINC01247 | 1.67669 | 1.73685 | 4.61555 | 0.000422783 | 0.015028174 | 0.25239 |
| CDC14C | 1.72583 | 0.72640 | 4.60561 | 0.000430749 | 0.015278751 | 0.22530 |
| C3orf33 | 1.52779 | 2.18721 | 4.60328 | 0.000432638 | 0.015313182 | 0.22584 |
| HTN1 | 1.58874 | 2.05029 | 4.60023 | 0.000435119 | 0.01534093 | 0.20277 |
| TYW5 | 1.13643 | 4.38182 | 4.61224 | 0.000435262 | 0.01534093 | -0.06839 |
| CENPK | 1.05333 | 3.16076 | 4.59737 | 0.000437462 | 0.015385922 | 0.10649 |
| AKAP5 | 1.45580 | 2.05251 | 4.59603 | 0.000438562 | 0.015392147 | 0.21169 |
| FOXG1-AS1 | 2.10470 | 0.00661 | 4.58711 | 0.000445978 | 0.015585527 | -0.11867 |
| ZNRF3-AS1 | 1.96198 | 1.76014 | 4.59688 | 0.000446584 | 0.015585527 | 0.20346 |
| ANKRD20A5P | 1.27445 | 2.95152 | 4.58535 | 0.000447461 | 0.015585527 | 0.14505 |
| CBWD3 | 1.34519 | 4.35425 | 4.59702 | 0.000447812 | 0.015585527 | -0.12801 |
| NAV3 | 1.38242 | 2.38563 | 4.58298 | 0.000449458 | 0.015610221 | 0.19032 |
| PCAT14 | 1.63701 | 1.93282 | 4.57728 | 0.0004543 | 0.015706252 | 0.18144 |
| LMO7-AS1 | 1.60089 | 0.35265 | 4.57666 | 0.000454828 | 0.015706252 | 0.14105 |
| FCAR | 1.47263 | 2.83906 | 4.58684 | 0.000455049 | 0.015706252 | 0.10632 |
| UNC80 | 1.15043 | 3.32716 | 4.57308 | 0.000457904 | 0.015750073 | 0.00783 |
| TEN1-CDK3 | 1.47314 | 1.38657 | 4.57273 | 0.000458208 | 0.015750073 | 0.17850 |
| WTIP | 1.19873 | 4.59551 | 4.58000 | 0.000462282 | 0.015857399 | -0.15833 |
| PPP1R3E | 1.06656 | 2.66037 | 4.56541 | 0.000464559 | 0.015902787 | 0.11157 |
| ZNF529-AS1 | 1.31865 | 2.72790 | 4.56275 | 0.000466889 | 0.015949796 | 0.11742 |
| STAG3L2 | 1.70954 | 3.33662 | 4.57113 | 0.00047002 | 0.016023923 | 0.01755 |
| LINC02618 | 1.73678 | 0.54262 | 4.55717 | 0.000471825 | 0.016052645 | 0.02872 |
| ZNF492 | 1.64587 | 1.18231 | 4.55326 | 0.000475311 | 0.016138308 | 0.14372 |
| LRRC36 | 1.57700 | 1.35137 | 4.54827 | 0.000479796 | 0.016232207 | 0.13626 |
| OSMR-AS1 | 2.24000 | -0.10719 | 4.54802 | 0.000480024 | 0.016232207 | -0.04556 |
| PRH1 | 1.92190 | 1.66516 | 4.54597 | 0.000481886 | 0.016262195 | 0.13198 |
| PMFBP1 | 1.54693 | 0.75958 | 4.54315 | 0.000484455 | 0.016315867 | 0.09098 |
| CYP2G1P | 2.02154 | 0.84563 | 4.54160 | 0.000485869 | 0.016330478 | 0.09521 |
| TMEM234 | 0.86010 | 4.32712 | 4.53877 | 0.000488473 | 0.016368283 | -0.18186 |
| LINC02826 | 1.68339 | 0.29673 | 4.53774 | 0.000489421 | 0.016368283 | 0.10998 |
| LINC00852 | 1.81096 | 0.97549 | 4.53718 | 0.000489939 | 0.016368283 | 0.09826 |
| TBXA2R | 1.49718 | 2.61669 | 4.54370 | 0.0004925 | 0.016402378 | 0.04550 |
| LINC02882 | 2.02510 | 0.96170 | 4.53356 | 0.000493295 | 0.016402378 | 0.03584 |
| LRIG2-DT | 1.47953 | 2.82234 | 4.54404 | 0.000494484 | 0.016402378 | 0.02416 |
| RAB40A | 1.82811 | 0.15241 | 4.53168 | 0.000495041 | 0.016402378 | 0.05518 |
| EAF1-AS1 | 1.66748 | 0.98894 | 4.53079 | 0.000495879 | 0.016402378 | 0.10048 |
| CLIC3 | 0.66093 | 3.75461 | 4.52885 | 0.000497691 | 0.016429713 | -0.16694 |
| LINC01232 | 1.14732 | 4.71559 | 4.53703 | 0.000501023 | 0.01650703 | -0.23725 |
| CCDC150 | 1.43818 | 1.20718 | 4.52308 | 0.000503141 | 0.016544096 | 0.09304 |
| F5 | 1.35044 | 2.46760 | 4.51892 | 0.000507109 | 0.016604831 | 0.01120 |
| OSGEP | 0.98511 | 3.52542 | 4.51854 | 0.000507467 | 0.016604831 | -0.08204 |
| XRCC2 | 1.49723 | 2.73483 | 4.52135 | 0.000507976 | 0.016604831 | 0.03193 |
| CASC15 | 1.36886 | 3.07391 | 4.52108 | 0.000511016 | 0.016671525 | -0.01965 |
| FGD5 | 1.66801 | 1.52272 | 4.51359 | 0.000512237 | 0.016675303 | 0.07240 |
| SUZ12P1 | 1.13441 | 3.63389 | 4.51999 | 0.000513132 | 0.016675303 | -0.14045 |
| TIAF1 | 1.59045 | 2.10241 | 4.51101 | 0.000514739 | 0.016694959 | 0.07187 |
| CCDC7 | 1.38091 | 2.54883 | 4.50487 | 0.000520738 | 0.016856757 | 0.02495 |
| CSMD1 | 1.18053 | 1.29920 | 4.49983 | 0.000525727 | 0.016985262 | 0.01095 |
| ADAMTS4 | 1.35049 | 2.83973 | 4.49663 | 0.00052891 | 0.01703649 | 0.01804 |
| BTBD16 | 3.06371 | -0.74223 | 4.49564 | 0.000529898 | 0.01703649 | -0.49173 |
| ESCO2 | 1.50171 | 2.72126 | 4.50408 | 0.000530378 | 0.01703649 | -0.02041 |
| CFHR2 | 1.99225 | 1.78701 | 4.50164 | 0.000535445 | 0.017155414 | 0.03421 |
| CYP4F26P | 1.16000 | 3.46021 | 4.50095 | 0.000536139 | 0.017155414 | -0.18814 |
| LNCARSR | 2.21079 | -0.29913 | 4.48727 | 0.000538354 | 0.017162023 | -0.14020 |
| KCNJ15 | 1.26307 | 5.70108 | 4.49870 | 0.000538404 | 0.017162023 | -0.37976 |
| ZNF37BP | 1.26525 | 3.96305 | 4.49724 | 0.000539889 | 0.01716818 | -0.26730 |
| ITGA9-AS1 | 1.38654 | 3.46721 | 4.49648 | 0.000540657 | 0.01716818 | -0.15801 |
| SNX18P3 | 1.23870 | 3.96590 | 4.49540 | 0.000541758 | 0.017170436 | -0.23476 |
| PCDHGB9P | 2.08511 | 0.34771 | 4.48092 | 0.000544854 | 0.017235796 | 0.00040 |
| CD99P1 | 1.34521 | 1.69065 | 4.47795 | 0.000547929 | 0.017300227 | 0.00524 |
| RRS1-AS1 | 1.47624 | 1.39028 | 4.46889 | 0.000557407 | 0.017566212 | -0.00071 |
| PSTK | 1.36874 | 1.57560 | 4.46672 | 0.000559698 | 0.017605136 | -0.00408 |
| TAGLN | 1.54149 | 3.07596 | 4.47144 | 0.000566392 | 0.017782157 | -0.07664 |
| LAMC3 | 2.52892 | -0.24690 | 4.44697 | 0.000581036 | 0.018207608 | -0.20367 |
| ZNF433-AS1 | 1.34850 | 2.35357 | 4.44447 | 0.000583801 | 0.018259925 | -0.07401 |
| DMC1 | 1.53736 | 1.53529 | 4.44253 | 0.000585951 | 0.018292866 | -0.04870 |
| LINC00174 | 1.10266 | 4.72035 | 4.44882 | 0.000591415 | 0.018428936 | -0.45590 |
| ENTPD1 | 1.28992 | 3.45686 | 4.44780 | 0.000592555 | 0.01843001 | -0.26336 |
| ZNF726 | 1.33626 | 2.87523 | 4.43469 | 0.000594724 | 0.018463026 | -0.14019 |
| ATAD3C | 2.26294 | 0.86116 | 4.43210 | 0.000597658 | 0.018519613 | -0.18762 |
| CYSLTR1 | 1.15712 | 4.39648 | 4.44056 | 0.000600691 | 0.01856333 | -0.43063 |
| LINC01136 | 2.26082 | -0.38453 | 4.42890 | 0.000601296 | 0.01856333 | -0.22131 |
| DYNAP | 2.94296 | 0.06556 | 4.42735 | 0.000603064 | 0.018583526 | -0.34417 |
| SLC27A5 | 1.22791 | 3.90234 | 4.43471 | 0.00060736 | 0.018681367 | -0.35774 |
| MMP25-AS1 | 1.61185 | 2.18300 | 4.41985 | 0.000611714 | 0.018780643 | -0.09385 |
| RPAIN | 1.16046 | 4.50149 | 4.42136 | 0.000622861 | 0.019087716 | -0.45183 |
| TMEM119 | 1.88765 | 0.03526 | 4.40810 | 0.000625522 | 0.019134091 | -0.19396 |
| MAILR | 1.28738 | 2.62056 | 4.40611 | 0.000627885 | 0.019171196 | -0.17192 |
| ZNF727 | 2.92777 | -0.56325 | 4.39885 | 0.000636611 | 0.019402085 | -0.74908 |
| LINC02444 | 1.38560 | 3.89559 | 4.40757 | 0.000639299 | 0.019448456 | -0.42119 |
| LINC00964 | 1.29533 | 2.20979 | 4.39286 | 0.000643894 | 0.019552551 | -0.19482 |
| MRO | 1.90844 | 1.29796 | 4.38908 | 0.000648538 | 0.019657787 | -0.13815 |
| SNHG4 | 1.17355 | 3.88151 | 4.39443 | 0.000655373 | 0.019828907 | -0.44189 |
| LINC01355 | 1.93424 | 2.58031 | 4.39090 | 0.000659769 | 0.019925736 | -0.15621 |
| OR6Y1 | 1.45558 | 1.77121 | 4.37190 | 0.000670098 | 0.020201102 | -0.16780 |
| TSEN2 | 0.97005 | 3.84763 | 4.36770 | 0.000675477 | 0.02032648 | -0.41868 |
| OR10AH1P | 1.70840 | 0.32132 | 4.36358 | 0.000680794 | 0.020411798 | -0.19689 |
| ZNF430 | 0.84739 | 4.16251 | 4.36164 | 0.000683316 | 0.020411798 | -0.50312 |
| RAB30-DT | 1.25198 | 3.47720 | 4.37202 | 0.000683766 | 0.020411798 | -0.37699 |
| ZSCAN30 | 0.95020 | 4.46359 | 4.37145 | 0.000684511 | 0.020411798 | -0.55013 |
| DUSP8P5 | 1.83210 | 0.40860 | 4.35935 | 0.000686301 | 0.020411798 | -0.19924 |
| CEACAM19 | 1.71257 | 1.51359 | 4.35919 | 0.000686511 | 0.020411798 | -0.20047 |
| MIR34AHG | 1.27787 | 4.18730 | 4.36962 | 0.000686883 | 0.020411798 | -0.52822 |
| LINC02301 | 1.47820 | 3.85386 | 4.36636 | 0.000691135 | 0.020501623 | -0.45936 |
| PTPRC | 2.67680 | -0.91258 | 4.35395 | 0.000693399 | 0.020532241 | -0.63560 |
| CAMK2A | 1.67869 | 1.44189 | 4.35166 | 0.000696432 | 0.020585498 | -0.20308 |
| NAIPP3 | 2.56539 | 1.31496 | 4.36107 | 0.000698102 | 0.020592198 | -0.22167 |
| ERVK3-1 | 0.85728 | 4.57860 | 4.34963 | 0.000699129 | 0.020592198 | -0.57423 |
| NPIPB12 | 1.42729 | 4.40991 | 4.35539 | 0.000705647 | 0.020747507 | -0.51491 |
| SLC38A6 | 0.98067 | 4.44197 | 4.34687 | 0.000711505 | 0.020877826 | -0.53901 |
| ZNF460-AS1 | 2.09097 | 0.36604 | 4.33963 | 0.000712584 | 0.020877826 | -0.30520 |
| IRF2-DT | 1.36834 | 2.14197 | 4.33742 | 0.000715601 | 0.020929455 | -0.25117 |
| DERPC | -1.05660 | 2.95383 | -4.33111 | 0.000724257 | 0.021145507 | -0.34234 |
| CYB5RL | 0.91589 | 4.24862 | 4.32861 | 0.000728744 | 0.021239319 | -0.57583 |
| ESRG | 1.29784 | 4.70444 | 4.33516 | 0.000733243 | 0.021303241 | -0.64286 |
| LENG8-AS1 | 1.28134 | 2.71149 | 4.32439 | 0.000733613 | 0.021303241 | -0.30709 |
| ODC1-DT | 1.74987 | 1.62616 | 4.32356 | 0.000734771 | 0.021303241 | -0.25478 |
| MIRLET7A1HG | 1.89147 | 2.66127 | 4.32637 | 0.000737102 | 0.021333727 | -0.25509 |
| AFDN-DT | 1.76836 | 0.71418 | 4.31896 | 0.000741248 | 0.021416542 | -0.26826 |
| TBILA | 1.21162 | 3.41845 | 4.32828 | 0.000742878 | 0.021426504 | -0.45513 |
| TDRKH-AS1 | 1.90220 | 0.92491 | 4.31490 | 0.000747015 | 0.021508624 | -0.29633 |
| SFTPD-AS1 | 1.54052 | 2.01795 | 4.31378 | 0.000748624 | 0.021517766 | -0.26942 |
| COL1A2 | 3.05663 | -0.05614 | 4.31219 | 0.000750889 | 0.021545728 | -0.78003 |
| UXT-AS1 | 1.69615 | 0.78336 | 4.30897 | 0.000755523 | 0.021635843 | -0.30086 |
| SPANXA2-OT1 | 1.89730 | 2.29426 | 4.30801 | 0.000756911 | 0.021635843 | -0.31670 |
| CCDC200 | 1.88996 | 0.57150 | 4.30699 | 0.00075839 | 0.021635843 | -0.38190 |
| ZNF491 | 1.13440 | 2.05824 | 4.30642 | 0.000759221 | 0.021635843 | -0.29922 |
| PCDHA10 | 1.47438 | 1.66022 | 4.30267 | 0.00076468 | 0.021684534 | -0.30150 |
| KIF5C | 1.87444 | 1.82989 | 4.30129 | 0.000766695 | 0.021684534 | -0.29008 |
| SYNE3 | 1.43880 | 1.82895 | 4.30112 | 0.000766946 | 0.021684534 | -0.30406 |
| ESYT3 | 1.59659 | 0.96688 | 4.30085 | 0.000767345 | 0.021684534 | -0.29243 |
| OAZ3 | 1.26946 | 1.70275 | 4.30079 | 0.000767433 | 0.021684534 | -0.30125 |
| TLL2 | 1.47362 | 0.84575 | 4.29704 | 0.00077295 | 0.021745928 | -0.30426 |
| ZNF439 | 1.16886 | 3.16582 | 4.29632 | 0.000774017 | 0.021745928 | -0.41209 |
| LINC02256 | 1.83258 | -1.08748 | 4.29539 | 0.000775382 | 0.021745928 | -0.46043 |
| WBP2NL | 1.60362 | 1.09240 | 4.29504 | 0.000775904 | 0.021745928 | -0.30263 |
| JMJD7-PLA2G4B | 1.61083 | 4.33845 | 4.30523 | 0.000776128 | 0.021745928 | -0.65940 |
| ZNF528-AS1 | 1.25044 | 2.38280 | 4.29078 | 0.000782247 | 0.021880598 | -0.36412 |
| SMIM36 | 2.60737 | -1.10011 | 4.28966 | 0.000783928 | 0.021890903 | -0.62137 |
| CTCFL | 1.85258 | -0.11190 | 4.28861 | 0.000785506 | 0.021898265 | -0.33280 |
| CYP2D7 | 2.21563 | 0.57954 | 4.28494 | 0.000791037 | 0.022015667 | -0.34142 |
| MGP | 1.75538 | 0.76647 | 4.27602 | 0.000804651 | 0.022357226 | -0.35577 |
| PIPOX | 2.07692 | 0.24919 | 4.27471 | 0.000806677 | 0.022376223 | -0.36329 |
| HCG25 | 1.30297 | 2.29590 | 4.27299 | 0.000809336 | 0.022412694 | -0.41814 |
| FEZ1 | 1.71782 | 1.86068 | 4.26944 | 0.000814848 | 0.022523688 | -0.34518 |
| C5orf64 | 1.44453 | 2.85402 | 4.26867 | 0.000816046 | 0.022523688 | -0.40187 |
| SKINT1L | 1.33146 | 4.50555 | 4.27292 | 0.000825322 | 0.022742052 | -0.80057 |
| MAGEA10 | 2.17424 | -0.07938 | 4.26172 | 0.00082698 | 0.022750136 | -0.43723 |
| DAND5 | 1.56379 | 0.92518 | 4.25927 | 0.000830873 | 0.022754392 | -0.36282 |
| FAM3D-AS1 | 1.58334 | 0.61167 | 4.25888 | 0.000831485 | 0.022754392 | -0.36943 |
| CFL1P1 | 1.42374 | 0.43933 | 4.25825 | 0.000832487 | 0.022754392 | -0.37270 |
| LINC01511 | 2.17444 | -0.73965 | 4.25819 | 0.000832594 | 0.022754392 | -0.53843 |
| PLCG1-AS1 | 1.84595 | 0.99855 | 4.25602 | 0.00083606 | 0.022811702 | -0.37263 |
| ZNF738 | 1.15993 | 2.91891 | 4.25416 | 0.000839046 | 0.022855785 | -0.48550 |
| NPIPB7 | 1.46001 | 1.76793 | 4.25107 | 0.000844016 | 0.022949136 | -0.37719 |
| TPM3P9 | 1.27433 | 2.53559 | 4.24967 | 0.000846281 | 0.022949136 | -0.42994 |
| PRDX6-AS1 | 1.65467 | 1.59767 | 4.24948 | 0.000846603 | 0.022949136 | -0.38913 |
| SNCA | -0.61996 | 3.67618 | -4.24734 | 0.000850071 | 0.023005723 | -0.72126 |
| TSTD3 | 1.09922 | 3.25758 | 4.24457 | 0.000854606 | 0.023090978 | -0.56495 |
| CRYZL2P | 0.73160 | 4.70153 | 4.24182 | 0.000859111 | 0.023151275 | -0.79075 |
| LINC00954 | 1.47350 | 1.48990 | 4.24151 | 0.000859626 | 0.023151275 | -0.39458 |
| CLCA4-AS1 | 1.57955 | -0.41900 | 4.24030 | 0.000861618 | 0.023151275 | -0.44779 |
| LRP5L | 1.54529 | 1.69377 | 4.23983 | 0.000862392 | 0.023151275 | -0.40810 |
| LINC00836 | 1.40246 | 0.97807 | 4.23056 | 0.000877864 | 0.023524558 | -0.41450 |
| ERICH6-AS1 | 1.00160 | 3.63307 | 4.22862 | 0.000881132 | 0.023524558 | -0.65734 |
| PARGP1 | 1.13068 | 3.38443 | 4.22823 | 0.000881801 | 0.023524558 | -0.59104 |
| ADAM32 | 1.39905 | 4.85077 | 4.23737 | 0.00088317 | 0.023524558 | -0.80577 |
| KCTD19 | 2.02813 | 0.27908 | 4.22731 | 0.000883353 | 0.023524558 | -0.44476 |
| ZNF487 | 0.88403 | 4.44730 | 4.22516 | 0.00089112 | 0.023656833 | -0.79281 |
| PI4KAP2 | 1.19404 | 3.17905 | 4.23265 | 0.000891158 | 0.023656833 | -0.63210 |
| LINC01795 | 1.94380 | -0.56485 | 4.21778 | 0.000899656 | 0.023844462 | -0.53674 |
| MEFV | 1.50955 | 3.02010 | 4.22474 | 0.000904718 | 0.02393736 | -0.57601 |
| ZNF66 | 0.97164 | 4.04945 | 4.21895 | 0.000906776 | 0.02393736 | -0.75509 |
| KLHL33 | 1.79967 | 1.25983 | 4.21209 | 0.000909539 | 0.02393736 | -0.49691 |
| ZNF662 | 1.17120 | 2.18781 | 4.21198 | 0.000909725 | 0.02393736 | -0.48409 |
| ANKRD36B | 0.93499 | 4.73326 | 4.22132 | 0.000910635 | 0.02393736 | -0.96301 |
| SLC26A8 | 1.23878 | 1.86900 | 4.21081 | 0.000911776 | 0.02393736 | -0.50025 |
| SLC14A1 | 2.62323 | -0.73764 | 4.20792 | 0.000916847 | 0.024032635 | -0.65413 |
| HIF1A-AS3 | 1.08159 | 4.66023 | 4.21195 | 0.000927075 | 0.024262593 | -0.92549 |
| LERFS | 1.55579 | 2.74242 | 4.20584 | 0.000937962 | 0.024509043 | -0.56058 |
| LINC01856 | 2.15389 | -0.21571 | 4.19228 | 0.0009448 | 0.024613167 | -0.63876 |
| PDE1C | 1.67480 | 1.53954 | 4.19156 | 0.000946116 | 0.024613167 | -0.48753 |
| RASAL2-AS1 | 1.39751 | 4.59022 | 4.20117 | 0.000946376 | 0.024613167 | -0.92542 |
| TMPRSS11BNL | 3.42196 | 0.36865 | 4.19704 | 0.000953872 | 0.024769463 | -0.60974 |
| LINC02101 | 2.43784 | -0.39911 | 4.17968 | 0.000967961 | 0.025081502 | -0.87533 |
| LINC01409 | 1.43211 | 2.30703 | 4.17918 | 0.000968897 | 0.025081502 | -0.50477 |
| OR56A1 | 2.00122 | 0.15998 | 4.17745 | 0.000972123 | 0.025125996 | -0.57186 |
| NCR3LG1 | 0.99152 | 3.65083 | 4.17330 | 0.000979926 | 0.025264592 | -0.74175 |
| LINC01694 | 1.52224 | 1.56364 | 4.17298 | 0.000980517 | 0.025264592 | -0.51300 |
| RBM14-RBM4 | 1.01156 | 2.66484 | 4.17189 | 0.000982581 | 0.025277617 | -0.59913 |
| LINC01087 | 1.56308 | 1.77196 | 4.17091 | 0.000984435 | 0.025277617 | -0.52384 |
| MCTS1 | 0.93226 | 4.68478 | 4.17955 | 0.000986308 | 0.025277617 | -0.92232 |
| P2RX5-TAX1BP3 | 1.64739 | 2.67411 | 4.17897 | 0.000987404 | 0.025277617 | -0.61856 |
| PHKG1 | 2.01029 | 0.77244 | 4.16827 | 0.000989695 | 0.025277617 | -0.52186 |
| PTBP2 | 0.93786 | 4.00543 | 4.17193 | 0.000991115 | 0.025277617 | -0.85892 |
| SNX18P8 | 1.51928 | 0.58341 | 4.16673 | 0.000992382 | 0.025277617 | -0.52667 |
| ZNF782 | 1.26472 | 2.97214 | 4.17141 | 0.00099326 | 0.025277617 | -0.66893 |
| KLF3-AS1 | 1.37765 | 3.25382 | 4.17514 | 0.000994668 | 0.025277617 | -0.72411 |
| SUCLG2-AS1 | 1.04569 | 3.30601 | 4.16350 | 0.000998561 | 0.025337905 | -0.73567 |
| LINC01910 | 2.05752 | -0.48141 | 4.16109 | 0.001003209 | 0.025409121 | -0.60489 |
| HTRA4 | 2.06028 | 0.96818 | 4.15911 | 0.00100704 | 0.025409121 | -0.67062 |
| LINC02057 | 1.28410 | 1.70528 | 4.15903 | 0.001007186 | 0.025409121 | -0.55906 |
| RPL4P6 | 2.45107 | -0.25650 | 4.15889 | 0.001007464 | 0.025409121 | -0.73125 |
| RHOF | 1.48603 | 3.14999 | 4.16321 | 0.001015172 | 0.025545149 | -0.61670 |
| PHBP19 | 2.66747 | -1.20442 | 4.15455 | 0.001015922 | 0.025545149 | -0.84998 |
| ZFP42 | 1.69459 | 1.24621 | 4.14708 | 0.001030635 | 0.025876078 | -0.55732 |
| EPHA10 | 1.49000 | 2.32025 | 4.15081 | 0.001038586 | 0.02601126 | -0.63517 |
| UGT1A1 | 1.59181 | 2.03413 | 4.15118 | 0.00103914 | 0.02601126 | -0.61355 |
| SLC26A4-AS1 | 1.03987 | 4.37465 | 4.15091 | 0.001041887 | 0.026040922 | -0.96409 |
| LINC00700 | 1.94997 | -0.29641 | 4.13274 | 0.001059496 | 0.0264414 | -0.75995 |
| RN7SL670P | 1.69071 | 1.41652 | 4.12706 | 0.001071155 | 0.026692417 | -0.59631 |
| POU5F1 | 1.66874 | 2.46236 | 4.13277 | 0.001078753 | 0.026841637 | -0.64706 |
| FBXO15 | 0.99694 | 4.87520 | 4.13126 | 0.001081885 | 0.026879436 | -1.04237 |
| BMS1P1 | 1.18448 | 4.84284 | 4.12773 | 0.00108923 | 0.027021645 | -1.09705 |
| NANOGP5 | 2.14737 | -0.67182 | 4.11630 | 0.00109361 | 0.027069265 | -0.75122 |
| SUGT1P2 | 2.10044 | -1.26663 | 4.11593 | 0.001094397 | 0.027069265 | -0.80913 |
| OPRM1 | 1.27781 | 1.95370 | 4.11363 | 0.00109927 | 0.027149515 | -0.64424 |
| CD180 | 1.45355 | 3.18259 | 4.12089 | 0.001103619 | 0.027216628 | -0.81985 |
| CBR3-AS1 | 1.16705 | 3.71511 | 4.11473 | 0.001116731 | 0.027493956 | -0.88183 |
| LINC00278 | 1.44582 | 2.05913 | 4.10479 | 0.001118163 | 0.027493956 | -0.66621 |
| LAMB4 | 1.52187 | 1.58054 | 4.09648 | 0.001136258 | 0.027897731 | -0.65196 |
| HTN3 | 2.21851 | -0.17467 | 4.09295 | 0.001144024 | 0.028047096 | -0.68458 |
| TSSK1B | 2.23183 | -0.09583 | 4.09087 | 0.001148625 | 0.028114569 | -0.84254 |
| KCTD9P4 | 1.57280 | -0.25507 | 4.09018 | 0.001150149 | 0.028114569 | -0.72820 |
| LRRFIP1P1 | 1.52593 | 0.20973 | 4.08138 | 0.001169865 | 0.028522165 | -0.69188 |
| PROX1-AS1 | 2.28076 | 0.35162 | 4.08121 | 0.001170245 | 0.028522165 | -0.88937 |
| RN7SL48P | 1.68834 | 0.27436 | 4.07420 | 0.001186197 | 0.028868737 | -0.69241 |
| OR7C1 | 1.57604 | 1.27765 | 4.07297 | 0.001189032 | 0.028895558 | -0.68629 |
| LINC01004 | 1.71104 | 0.88551 | 4.07030 | 0.001195187 | 0.028986302 | -0.69432 |
| ADCY10P1 | 1.27003 | 1.41988 | 4.06946 | 0.001197125 | 0.028986302 | -0.70142 |
| BDNF-AS | 0.94910 | 2.90289 | 4.06909 | 0.001197982 | 0.028986302 | -0.84317 |
| ZSCAN4 | 1.35311 | 1.36893 | 4.06820 | 0.001200038 | 0.02899397 | -0.69570 |
| CSMD3 | 2.49268 | -0.18001 | 4.06538 | 0.001206604 | 0.029110417 | -0.93355 |
| CTSK | 0.86338 | 2.85770 | 4.06427 | 0.001209198 | 0.029130831 | -0.87401 |
| GRIK2 | 1.03557 | 3.10024 | 4.05321 | 0.001235321 | 0.02968124 | -0.84195 |
| OR6J1 | 1.51177 | 1.34460 | 4.05309 | 0.001235606 | 0.02968124 | -0.73064 |
| PDXDC2P | 1.12386 | 1.52215 | 4.05104 | 0.001240524 | 0.029756511 | -0.73903 |
| CCT6P3 | 1.28204 | 2.51296 | 4.04919 | 0.001244977 | 0.029756519 | -0.82232 |
| ZRANB2-AS2 | 1.25607 | 1.11802 | 4.04834 | 0.001247018 | 0.029756519 | -0.73079 |
| TCAF2 | 1.48155 | 3.00820 | 4.05667 | 0.001248545 | 0.029756519 | -0.87056 |
| DELEC1 | 1.92266 | -0.62966 | 4.04724 | 0.00124968 | 0.029756519 | -0.95204 |
| RASGRF2-AS1 | 1.85583 | 0.59487 | 4.04672 | 0.001250927 | 0.029756519 | -0.79183 |
| SCOC-AS1 | 1.57298 | 0.48024 | 4.04660 | 0.001251234 | 0.029756519 | -0.73436 |
| STX17-AS1 | 1.02397 | 4.55945 | 4.05421 | 0.001254481 | 0.029791252 | -1.19130 |
| PTCSC2 | 1.32448 | 1.57334 | 4.04375 | 0.001258134 | 0.029835487 | -0.75893 |
| PMS2P4 | 1.79777 | 0.39808 | 4.04168 | 0.001263185 | 0.029912729 | -0.75733 |
| EXTL3-AS1 | 1.29656 | 2.30953 | 4.04067 | 0.001265665 | 0.029921692 | -0.78670 |
| BLOC1S6P1 | 1.50346 | 0.31380 | 4.04006 | 0.001267153 | 0.029921692 | -0.75578 |
| SEPTIN14 | 1.37616 | 1.12717 | 4.03556 | 0.001278252 | 0.03014107 | -0.75266 |
| TTTY14 | 1.16938 | 2.25064 | 4.03243 | 0.001286018 | 0.030281359 | -0.79828 |
| HHIPL1 | 1.97760 | -0.04279 | 4.02901 | 0.00129456 | 0.030439516 | -0.84702 |
| DLC1 | 2.25641 | -0.05107 | 4.02827 | 0.001296413 | 0.030440141 | -0.79942 |
| TTC6 | 0.97685 | 4.98417 | 4.03612 | 0.001298922 | 0.030456151 | -1.26318 |
| FAM71F2 | 1.53222 | 2.60800 | 4.02257 | 0.001313587 | 0.030756743 | -0.81695 |
| SPICE1 | 0.84658 | 4.80149 | 4.02574 | 0.001317569 | 0.030806718 | -1.20962 |
| ANKRD18B | 0.96101 | 4.28518 | 4.02747 | 0.001320763 | 0.030826937 | -1.18827 |
| PCAT19 | 1.76521 | 0.83479 | 4.01765 | 0.00132334 | 0.030826937 | -0.78166 |
| GPR137C | 1.38673 | 0.67274 | 4.01715 | 0.001324633 | 0.030826937 | -0.78225 |
| TPH1 | 1.02679 | 1.89207 | 4.01640 | 0.001326558 | 0.030826937 | -0.86486 |
| CASC2 | 0.87999 | 5.62850 | 4.02476 | 0.001327679 | 0.030826937 | -1.28581 |
| IQCM | 2.19387 | 0.28922 | 4.01201 | 0.001337875 | 0.030948876 | -0.87945 |
| LIAS | 1.04981 | 4.16681 | 4.02073 | 0.001338029 | 0.030948876 | -1.15676 |
| RASGRP4 | 1.43678 | 1.24579 | 4.01177 | 0.0013385 | 0.030948876 | -0.79778 |
| LINC00996 | 1.67539 | 0.41800 | 4.01098 | 0.00134055 | 0.030953344 | -0.80007 |
| CAPRIN2 | 0.89044 | 4.15144 | 4.01388 | 0.001345206 | 0.030992288 | -1.17336 |
| LINC00910 | 1.08045 | 3.73319 | 4.01746 | 0.001346476 | 0.030992288 | -1.11020 |
| ETF1P2 | 2.64639 | -0.26412 | 4.00819 | 0.001347814 | 0.030992288 | -0.97708 |
| LOH12CR2 | 1.91056 | 0.35707 | 4.00706 | 0.00135079 | 0.031017929 | -0.83376 |
| DPY19L1P1 | 1.10486 | 1.75796 | 4.00504 | 0.001356077 | 0.031095399 | -0.81683 |
| TNRC18P1 | 2.07316 | 0.32352 | 4.00395 | 0.001358948 | 0.031095399 | -0.82769 |
| RALGAPA1P1 | 1.73377 | 2.18420 | 4.00364 | 0.001359759 | 0.031095399 | -0.80870 |
| DCT | 1.92559 | -0.63020 | 3.99975 | 0.001370067 | 0.031256284 | -0.84617 |
| CHKB | 0.78742 | 4.50486 | 3.99957 | 0.001370544 | 0.031256284 | -1.22923 |
| OR7E7P | 1.90144 | 0.22477 | 3.99863 | 0.001373032 | 0.03127025 | -0.90057 |
| PCAT1 | 1.21239 | 3.36979 | 3.99582 | 0.001380543 | 0.031396456 | -0.92878 |
| GBAP1 | 1.22393 | 0.96385 | 3.99459 | 0.001383837 | 0.031396456 | -0.82931 |
| TPT1-AS1 | 1.15765 | 4.28193 | 4.00312 | 0.001384224 | 0.031396456 | -1.23757 |
| RN7SL648P | 2.52962 | -0.82229 | 3.99231 | 0.001389958 | 0.03148369 | -1.23423 |
| PCDHGA8 | 1.46886 | 2.01113 | 3.99178 | 0.001397076 | 0.031601963 | -0.88945 |
| ZNF333 | 0.87075 | 3.28749 | 3.98683 | 0.001404812 | 0.031733914 | -1.04460 |
| FAM221A | 1.16052 | 4.73668 | 3.99312 | 0.001411203 | 0.031835131 | -1.26996 |
| SLC25A25-AS1 | 1.14718 | 4.41678 | 3.99225 | 0.001413559 | 0.03184519 | -1.23864 |
| LINC00507 | 1.88166 | 1.39152 | 3.98005 | 0.001423415 | 0.032023959 | -0.84693 |
| FBXL19-AS1 | 1.31664 | 2.06260 | 3.97899 | 0.001426327 | 0.03204623 | -0.88879 |
| XIST | 1.12184 | 6.73360 | 3.98174 | 0.001431443 | 0.032117879 | -1.06922 |
| PKD1L2 | 1.38140 | 1.77072 | 3.97493 | 0.001437626 | 0.032213267 | -0.89036 |
| GAPLINC | 2.75317 | -0.95602 | 3.97379 | 0.001440813 | 0.032224645 | -1.17989 |
| MOBP | 1.65548 | 0.43287 | 3.97333 | 0.001442075 | 0.032224645 | -0.87836 |
| PTGES3L | 1.33798 | 0.98562 | 3.97267 | 0.001443933 | 0.032224645 | -0.85992 |
| TM4SF19 | 1.39073 | 1.40382 | 3.97160 | 0.001446944 | 0.032247844 | -0.87824 |
| RPS15AP10 | 2.76996 | -1.13081 | 3.97077 | 0.001449262 | 0.032247844 | -1.35320 |
| SLC2A1-AS1 | 1.77244 | 1.16005 | 3.96934 | 0.001453303 | 0.032247844 | -0.87267 |
| C4BPB | 1.51496 | 1.53375 | 3.96906 | 0.001454076 | 0.032247844 | -0.89145 |
| SLC23A3 | 1.89167 | 0.39653 | 3.96886 | 0.001454645 | 0.032247844 | -0.88013 |
| MOGAT3 | 2.42175 | -0.15957 | 3.95768 | 0.001486578 | 0.032897278 | -0.98909 |
| MYLK3 | 1.21667 | 3.89483 | 3.96564 | 0.001488087 | 0.032897278 | -1.26118 |
| LINC02517 | 1.25565 | 3.16676 | 3.96478 | 0.001489859 | 0.032897278 | -1.05452 |
| ZNRD2-AS1 | 1.97376 | 0.13132 | 3.95455 | 0.001495625 | 0.032980899 | -0.91650 |
| KIAA1671-AS1 | 1.50504 | 0.33894 | 3.94935 | 0.001510816 | 0.033271884 | -0.90349 |
| LINC01115 | 1.64791 | -0.86127 | 3.93518 | 0.001552988 | 0.0341555 | -1.01219 |
| TAS2R13 | 2.55094 | -0.18224 | 3.93356 | 0.001557896 | 0.034218303 | -1.08813 |
| ZNF891 | 1.03462 | 4.51154 | 3.94063 | 0.001561789 | 0.034258657 | -1.40427 |
| KLF3P1 | 2.41953 | -1.07084 | 3.93157 | 0.001563916 | 0.034260235 | -1.06750 |
| POLN | 1.19850 | 2.52300 | 3.92481 | 0.001584607 | 0.034638223 | -1.04348 |
| PRR4 | 2.34727 | -0.94176 | 3.92416 | 0.001586601 | 0.034638223 | -1.08252 |
| LINC01546 | 2.05265 | 0.38608 | 3.92390 | 0.001587403 | 0.034638223 | -0.95817 |
| FAM221B | 1.34388 | 0.89794 | 3.92311 | 0.001589868 | 0.034646645 | -0.94722 |
| RAMP2-AS1 | 1.30043 | 2.21750 | 3.92152 | 0.001594793 | 0.034677341 | -1.04524 |
| ATP6V0CP4 | 2.92168 | -0.81806 | 3.92131 | 0.001595436 | 0.034677341 | -1.26403 |
| MEIOC | 2.24831 | -0.43054 | 3.91979 | 0.001600149 | 0.034734484 | -1.05606 |
| ARHGAP23P1 | 3.36227 | -1.22248 | 3.91641 | 0.001610708 | 0.034918231 | -1.86493 |
| ATP1B1P1 | 3.03717 | -1.12032 | 3.91567 | 0.001613029 | 0.034923133 | -1.69870 |
| CACNG8 | 1.65183 | 1.28782 | 3.91366 | 0.001619344 | 0.035014365 | -0.96220 |
| AKR1C8P | 3.54206 | -0.86260 | 3.91176 | 0.00162535 | 0.035098726 | -1.57697 |
| CD300LG | 1.74707 | 0.93010 | 3.90981 | 0.001631511 | 0.035186181 | -0.97288 |
| LINC00862 | 2.05517 | -0.07729 | 3.90803 | 0.001637173 | 0.035262683 | -1.13085 |
| ZSCAN16-AS1 | 1.42715 | 0.89812 | 3.90705 | 0.001640315 | 0.035284765 | -0.97806 |
| ASAP1-IT2 | 1.99670 | -0.48458 | 3.90542 | 0.001645522 | 0.035343049 | -1.01014 |
| IMPG2 | 1.15259 | 2.69131 | 3.90487 | 0.001647265 | 0.035343049 | -1.07235 |
| DLL1 | -0.53571 | 3.86081 | -3.90196 | 0.001656637 | 0.035441572 | -1.42259 |
| ACP6 | 0.94467 | 4.63523 | 3.90966 | 0.00165823 | 0.035441572 | -1.43876 |
| TIMP1 | -0.69420 | 5.24462 | -3.90146 | 0.001658234 | 0.035441572 | -1.50473 |
| LINC02453 | 1.91063 | 0.40889 | 3.89440 | 0.001681179 | 0.035868786 | -1.01966 |
| CHMP4BP1 | 1.79022 | -0.18059 | 3.89399 | 0.001682526 | 0.035868786 | -1.08047 |
| SHOX | 1.25686 | 2.59339 | 3.89557 | 0.001693587 | 0.036005413 | -1.15317 |
| FAM66D | 1.03509 | 2.19340 | 3.89028 | 0.001694736 | 0.036005413 | -1.09862 |
| RNF207 | 1.12399 | 4.60288 | 3.89821 | 0.001695414 | 0.036005413 | -1.54625 |
| NR5A2 | 1.52346 | 1.63217 | 3.88917 | 0.001698397 | 0.036022874 | -1.02265 |
| LINC00565 | 1.52674 | -0.11216 | 3.88731 | 0.001704544 | 0.036069706 | -1.02564 |
| GOLGA2P5 | 1.05867 | 5.06935 | 3.89532 | 0.001704932 | 0.036069706 | -1.59008 |
| PRAL | 1.50147 | 2.14580 | 3.88904 | 0.001715172 | 0.036240342 | -1.07811 |
| LMCD1-AS1 | 1.28440 | 1.41255 | 3.88271 | 0.001719896 | 0.036294151 | -1.03627 |
| CAPN10-DT | 1.68220 | 0.82839 | 3.87986 | 0.001729469 | 0.036450025 | -1.02299 |
| OR1I1 | 1.54084 | 2.26828 | 3.87377 | 0.001750096 | 0.036827914 | -1.06154 |
| GPATCH2 | 0.48414 | 4.87758 | 3.87327 | 0.001751817 | 0.036827914 | -1.51171 |
| MTCO1P11 | 1.81779 | -0.28601 | 3.86878 | 0.001767198 | 0.037104477 | -1.05314 |
| TRABD2A | 1.26583 | 2.23640 | 3.86766 | 0.00177105 | 0.037138592 | -1.10558 |
| COQ6 | 0.73358 | 4.42001 | 3.86691 | 0.001773632 | 0.037146014 | -1.45712 |
| SERPINA10 | 1.76843 | 0.99579 | 3.86124 | 0.001793362 | 0.037512093 | -1.05761 |
| FGF14-IT1 | 1.78307 | 0.47546 | 3.85975 | 0.001798555 | 0.037540394 | -1.07291 |
| ZDHHC11B | 1.01877 | 2.73567 | 3.85956 | 0.001799219 | 0.037540394 | -1.18696 |
| PRICKLE2-AS3 | 1.41974 | 1.64727 | 3.85665 | 0.001809454 | 0.037706764 | -1.09958 |
| LINC01322 | 1.48203 | 2.43818 | 3.85781 | 0.001817343 | 0.037823876 | -1.12601 |
| PIF1 | 1.42344 | 1.12131 | 3.85219 | 0.001825261 | 0.037929111 | -1.07155 |
| TMED2-DT | 1.23808 | 3.89298 | 3.85966 | 0.00182695 | 0.037929111 | -1.44938 |
| FOXP1-IT1 | 1.77060 | 1.72766 | 3.85031 | 0.001831953 | 0.037985674 | -1.07406 |
| SUGT1P3 | 1.55873 | 0.67278 | 3.84903 | 0.001836552 | 0.038033738 | -1.07264 |
| C5AR2 | 1.34610 | 3.57576 | 3.85519 | 0.001842843 | 0.038073606 | -1.29643 |
| GABRB3 | 1.26344 | 2.16070 | 3.84683 | 0.001844441 | 0.038073606 | -1.09774 |
| ZNF589 | 0.83687 | 3.72308 | 3.84623 | 0.001846593 | 0.038073606 | -1.46456 |
| TVP23C | 1.30567 | 4.98825 | 3.85386 | 0.001847612 | 0.038073606 | -1.46137 |
| SYN3 | 1.52893 | 1.78610 | 3.84425 | 0.001853733 | 0.038084259 | -1.08167 |
| GVQW3 | 1.00785 | 3.62749 | 3.85199 | 0.001854342 | 0.038084259 | -1.41802 |
| LDLRAD4 | 0.90248 | 5.90180 | 3.85123 | 0.001857088 | 0.038084259 | -1.60697 |
| SCIMP | 1.52607 | 1.49560 | 3.84328 | 0.001857267 | 0.038084259 | -1.10259 |
| SEC14L4 | 1.49175 | 1.23440 | 3.84188 | 0.001862325 | 0.038125708 | -1.08645 |
| NAP1L4P1 | 1.81070 | -0.04879 | 3.84146 | 0.001863862 | 0.038125708 | -1.10767 |
| LINC00621 | 1.12690 | 3.24807 | 3.84474 | 0.001876812 | 0.038305179 | -1.32471 |
| MAPK10 | 0.99131 | 7.33636 | 3.84567 | 0.001877232 | 0.038305179 | -1.60492 |
| LINC00501 | 1.73361 | 0.12079 | 3.83606 | 0.001883598 | 0.038334388 | -1.10536 |
| PLAC4 | 0.60022 | 3.92375 | 3.83587 | 0.001884273 | 0.038334388 | -1.45680 |
| FXN | 0.64554 | 3.63755 | 3.83498 | 0.001887562 | 0.038334388 | -1.40303 |
| LINC01344 | 1.38430 | 2.77989 | 3.83612 | 0.001887861 | 0.038334388 | -1.18125 |
| IFI6 | 1.68462 | 3.78203 | 3.83951 | 0.001899805 | 0.038529989 | -1.27112 |
| STAU2-AS1 | 1.34758 | 0.88682 | 3.83081 | 0.001902979 | 0.038547457 | -1.10400 |
| G6PC | 1.68627 | 0.03261 | 3.82465 | 0.001926002 | 0.038966488 | -1.14639 |
| CREG2 | 1.76943 | 2.57612 | 3.82787 | 0.00194323 | 0.039229196 | -1.17352 |
| LINC01505 | 1.87240 | 0.03900 | 3.81969 | 0.00194471 | 0.039229196 | -1.13983 |
| RASL10B | 1.33929 | 3.11358 | 3.82609 | 0.001946047 | 0.039229196 | -1.25293 |
| LINC02071 | 2.58198 | -0.02105 | 3.81998 | 0.001950799 | 0.039277499 | -1.18970 |
| TMEM45A | 1.08724 | 8.07798 | 3.82328 | 0.001960608 | 0.039427375 | -1.59096 |
| DLEU1 | 0.78031 | 4.35229 | 3.81056 | 0.001979672 | 0.039753475 | -1.56556 |
| CA14 | 1.52801 | 0.92322 | 3.81007 | 0.001981593 | 0.039753475 | -1.13985 |
| ZNF577 | 0.83816 | 3.12564 | 3.80830 | 0.001988455 | 0.039824485 | -1.36947 |
| GSDMB | 1.05859 | 4.40360 | 3.81550 | 0.001990464 | 0.039824485 | -1.57665 |
| OR4F17 | 0.98632 | 4.41682 | 3.81483 | 0.00199305 | 0.039824485 | -1.63813 |
| NSUN5P1 | 1.00561 | 4.66971 | 3.81441 | 0.001994688 | 0.039824485 | -1.70493 |
| ANKAR | 1.19052 | 2.36591 | 3.80464 | 0.002002696 | 0.039936537 | -1.23656 |
| TMEM30BP1 | 1.79282 | -0.18022 | 3.80244 | 0.00201133 | 0.040059556 | -1.18135 |
| WARS2-AS1 | 0.86163 | 4.89268 | 3.80901 | 0.002015736 | 0.040059556 | -1.68457 |
| CSPG4P12 | 1.68285 | 2.22498 | 3.80123 | 0.002016074 | 0.040059556 | -1.15935 |
| CCDC26 | 1.11534 | 3.69791 | 3.80439 | 0.002033925 | 0.040366154 | -1.42135 |
| PCDHB1 | 1.30964 | 2.75365 | 3.80225 | 0.002042383 | 0.040485813 | -1.33568 |
| TRIM72 | 1.44564 | 2.74501 | 3.80157 | 0.002045083 | 0.040491179 | -1.33379 |
| BMS1P10 | 2.43161 | 0.00161 | 3.79096 | 0.002056921 | 0.040677259 | -1.24503 |
| NPAS1 | -1.17918 | 0.69752 | -3.79014 | 0.00206022 | 0.040694223 | -1.17504 |
| ALOX15P1 | 1.11466 | 1.61967 | 3.78627 | 0.00207585 | 0.040954427 | -1.23068 |
| FHIT | 1.07924 | 2.82889 | 3.78434 | 0.002083712 | 0.041028704 | -1.35507 |
| SPTB | 1.36425 | 2.23286 | 3.78820 | 0.002084537 | 0.041028704 | -1.28799 |
| KLK7 | 1.69851 | 1.34774 | 3.78119 | 0.002096553 | 0.041191731 | -1.19300 |
| LINC00937 | 1.25498 | 1.88114 | 3.78090 | 0.002097761 | 0.041191731 | -1.26116 |
| CNIH3-AS2 | 1.85249 | 0.74685 | 3.77979 | 0.002102331 | 0.041232892 | -1.19784 |
| DISC1-IT1 | 1.50159 | 1.27494 | 3.77740 | 0.002112141 | 0.041376617 | -1.19870 |
| LINC02649 | 1.47540 | 1.62400 | 3.77612 | 0.002117432 | 0.041398554 | -1.19905 |
| CFAP92 | 0.83995 | 4.75593 | 3.78347 | 0.002118342 | 0.041398554 | -1.72154 |
| PTPN20 | 1.17541 | 1.91062 | 3.77533 | 0.002120711 | 0.041398554 | -1.23370 |
| SEC24B-AS1 | 1.25107 | 1.70624 | 3.77322 | 0.002129484 | 0.041521196 | -1.23125 |
| ST3GAL5 | 0.84228 | 4.56021 | 3.77916 | 0.002132017 | 0.041522027 | -1.64825 |
| HDAC1P1 | 2.00984 | -0.74827 | 3.76796 | 0.002151507 | 0.041817937 | -1.25707 |
| LINC02388 | 1.52286 | 1.01838 | 3.76778 | 0.002152228 | 0.041817937 | -1.21301 |
| LINC02371 | 1.88786 | -0.00548 | 3.76327 | 0.002171303 | 0.042139448 | -1.23201 |
| CYP2B7P | 0.79505 | 7.93984 | 3.76927 | 0.002175157 | 0.042161197 | -1.71980 |
| RNPC3 | 1.06179 | 4.44917 | 3.76912 | 0.002178311 | 0.042161197 | -1.68965 |
| ATAD3B | 0.92769 | 4.26818 | 3.76872 | 0.00218001 | 0.042161197 | -1.66691 |
| C4B | 0.86648 | 3.68501 | 3.75667 | 0.00219952 | 0.042486722 | -1.57558 |
| RAD17P1 | 1.45902 | 0.88478 | 3.75537 | 0.00220511 | 0.042486722 | -1.23475 |
| LINC01748 | 2.27626 | -0.16878 | 3.75526 | 0.002205586 | 0.042486722 | -1.40283 |
| LINC02395 | 1.37316 | 1.90567 | 3.75473 | 0.002207886 | 0.042486722 | -1.29170 |
| ARGFX | 1.84938 | 1.68955 | 3.75415 | 0.002210373 | 0.042486722 | -1.23604 |
| DHRS2 | 1.72081 | -0.50271 | 3.75374 | 0.002212133 | 0.042486722 | -1.27318 |
| HDAC8 | 1.18722 | 5.22108 | 3.75836 | 0.002224384 | 0.042641925 | -1.78939 |
| PARVB | 1.14450 | 3.16243 | 3.75340 | 0.00222533 | 0.042641925 | -1.43658 |
| BORCS8 | 1.06375 | 4.15311 | 3.75531 | 0.00223767 | 0.04282916 | -1.62938 |
| TAS2R64P | 2.07134 | 0.08880 | 3.74657 | 0.002243391 | 0.042839306 | -1.37078 |
| SH3GL1P1 | 2.68399 | 0.03102 | 3.74613 | 0.002245313 | 0.042839306 | -1.41045 |
| SETD4 | 0.90809 | 4.32177 | 3.75342 | 0.002245909 | 0.042839306 | -1.68849 |
| ZNF814 | 1.02077 | 5.12444 | 3.75083 | 0.002257237 | 0.043006164 | -1.81424 |
| GUSBP9 | 1.70598 | 0.54284 | 3.74237 | 0.002261882 | 0.04304547 | -1.25802 |
| NPHP3 | 0.90173 | 4.48967 | 3.74883 | 0.002266066 | 0.043075923 | -1.78276 |
| ZNF567 | 0.79375 | 3.53591 | 3.74027 | 0.002271196 | 0.043124263 | -1.54337 |
| CYB5AP2 | 1.23447 | 1.11328 | 3.73818 | 0.002280534 | 0.043252314 | -1.27613 |
| CEP57L1 | 0.78725 | 4.73648 | 3.74326 | 0.002290769 | 0.043397048 | -1.78796 |
| GGT5 | 1.65652 | 0.20875 | 3.73415 | 0.002298577 | 0.043495545 | -1.28430 |
| ADSL | 0.83793 | 4.66425 | 3.73934 | 0.002308333 | 0.043630642 | -1.77046 |
| TAS2R3 | 2.02337 | -0.38054 | 3.72883 | 0.002322617 | 0.043850899 | -1.32997 |
| APTX | 0.70659 | 5.19286 | 3.72967 | 0.002329919 | 0.043939001 | -1.83703 |
| PLA2G4B | 1.56293 | 2.44576 | 3.73099 | 0.002346164 | 0.044195373 | -1.50093 |
| ZNF418 | 0.99669 | 3.00432 | 3.72207 | 0.002353583 | 0.04425358 | -1.47891 |
| ARLNC1 | 2.65365 | -0.74474 | 3.72185 | 0.002354563 | 0.04425358 | -1.76924 |
| LINC00672 | 1.53465 | 0.99602 | 3.71918 | 0.002366932 | 0.044435952 | -1.29745 |
| PCP4L1 | -0.65079 | 4.11136 | -3.71575 | 0.002382878 | 0.044635529 | -1.80035 |
| CACNG5 | 1.58499 | 1.29909 | 3.71574 | 0.002382918 | 0.044635529 | -1.32423 |
| FBXW4P1 | 1.40655 | 0.74787 | 3.71053 | 0.002407332 | 0.045042245 | -1.31690 |
| GSN-AS1 | 2.22122 | -0.68973 | 3.70924 | 0.00241345 | 0.045106078 | -1.34968 |
| LINC01151 | 1.92602 | 0.89412 | 3.71453 | 0.002422649 | 0.045227306 | -1.32279 |
| ALMS1-IT1 | 1.55768 | 0.51125 | 3.70614 | 0.002428155 | 0.045279378 | -1.31848 |
| LINC01203 | 2.08685 | -0.20458 | 3.70220 | 0.002446949 | 0.045578865 | -1.35551 |
| RPSAP36 | 1.49404 | -0.08614 | 3.69949 | 0.002459967 | 0.045770216 | -1.33612 |
| STX18-AS1 | 1.10124 | 4.88570 | 3.70398 | 0.002472971 | 0.045960862 | -1.91707 |
| NTRK3 | 2.41988 | -0.51032 | 3.69551 | 0.002479217 | 0.046025651 | -1.49760 |
| AFG3L1P | 1.05396 | 3.06787 | 3.69251 | 0.002493826 | 0.046245359 | -1.53746 |
| HELLS | 0.68196 | 4.72710 | 3.69173 | 0.002497629 | 0.046264414 | -1.87201 |
| RNF7P1 | 1.49625 | 1.46466 | 3.69062 | 0.002503102 | 0.046314328 | -1.34568 |
| CXCL2 | -1.09946 | 2.63255 | -3.69113 | 0.002509124 | 0.046369241 | -1.56361 |
| CHRD | 1.99387 | 0.21470 | 3.68848 | 0.002513618 | 0.046369241 | -1.36845 |
| ZNF248 | 0.75994 | 4.12516 | 3.68832 | 0.002514414 | 0.046369241 | -1.80459 |
| LINC01970 | 1.25927 | 1.66571 | 3.68460 | 0.00253279 | 0.046656515 | -1.41060 |
| MICA | 1.08363 | 3.21739 | 3.68500 | 0.002538813 | 0.046715841 | -1.57966 |
| ETS1 | 0.53823 | 5.09011 | 3.68233 | 0.002544092 | 0.046761357 | -1.87800 |
| H3P37 | 1.66644 | 1.10765 | 3.67753 | 0.002568118 | 0.047117662 | -1.37887 |
| CNR2 | 1.34297 | 2.04659 | 3.67821 | 0.002569129 | 0.047117662 | -1.46481 |
| TAT | 1.76887 | 0.40418 | 3.67589 | 0.002576384 | 0.047198792 | -1.37366 |
| TBC1D3B | 1.54963 | 1.02820 | 3.67481 | 0.002581834 | 0.047246711 | -1.37933 |
| CEP44 | 0.59921 | 5.23258 | 3.67233 | 0.002594437 | 0.047425288 | -1.96071 |
| ZNF700 | 0.80661 | 3.80982 | 3.67111 | 0.002600642 | 0.047486637 | -1.74181 |
| SLC5A12 | 2.52133 | -0.13716 | 3.66872 | 0.002612875 | 0.047657804 | -1.44697 |
| LRRC70 | 2.31313 | -0.59413 | 3.66689 | 0.002622267 | 0.047776848 | -1.51837 |
| KCNC4 | 0.96700 | 4.42739 | 3.67285 | 0.002627815 | 0.047787029 | -1.92224 |
| ZNF90 | 1.13902 | 3.86879 | 3.67270 | 0.002628559 | 0.047787029 | -1.81741 |
| CBWD5 | 1.01053 | 4.69193 | 3.67096 | 0.002637519 | 0.047897684 | -1.94277 |
| SOCS5P4 | 2.11556 | -0.40142 | 3.66037 | 0.002655973 | 0.048157766 | -1.43940 |
| B3GALT5-AS1 | 1.01928 | 2.44587 | 3.65992 | 0.002658328 | 0.048157766 | -1.47152 |
| HYPK | 1.11549 | 3.50088 | 3.66651 | 0.002660506 | 0.048157766 | -1.68141 |
| LCN15 | 1.95390 | 0.08039 | 3.65776 | 0.002669586 | 0.048269703 | -1.54153 |
| LINC01106 | 1.22189 | 2.15756 | 3.65442 | 0.002687146 | 0.048493451 | -1.50748 |
| DTX2P1 | 1.09029 | 2.99183 | 3.66128 | 0.002687778 | 0.048493451 | -1.70035 |
| FAM106A | 1.04399 | 2.48852 | 3.65296 | 0.002694845 | 0.048528087 | -1.52596 |
| TTLL9 | 0.76745 | 4.93092 | 3.65981 | 0.00269552 | 0.048528087 | -2.00588 |
| LINC00893 | 1.13366 | 2.89515 | 3.65167 | 0.002701698 | 0.048556154 | -1.51937 |
| PCDHA4 | 1.31779 | 4.66153 | 3.65796 | 0.002705284 | 0.048556154 | -1.98410 |
| SIRPB1 | 1.31384 | 1.39749 | 3.65006 | 0.002710225 | 0.048556154 | -1.42900 |
| BCL2L15 | 0.86537 | 3.63814 | 3.64979 | 0.002711676 | 0.048556154 | -1.74178 |
| FFAR2 | 2.02955 | -0.54150 | 3.64936 | 0.002713969 | 0.048556154 | -1.46781 |
| RELA-DT | 1.15670 | 2.31837 | 3.64906 | 0.002715533 | 0.048556154 | -1.54904 |
| LINC00702 | 2.42748 | 0.56276 | 3.64866 | 0.002717705 | 0.048556154 | -1.51133 |
| FRMD6-AS1 | 1.76109 | -0.36113 | 3.64798 | 0.002721294 | 0.048556154 | -1.44729 |
| CASR | 1.13539 | 2.08692 | 3.64761 | 0.002723292 | 0.048556154 | -1.50321 |
| FAM66B | 1.20298 | 2.52053 | 3.64748 | 0.002736626 | 0.048717104 | -1.58678 |
| CCDC159 | 0.74691 | 3.28036 | 3.64483 | 0.002738164 | 0.048717104 | -1.69284 |
| GPRASP1 | 1.33247 | 1.73685 | 3.64382 | 0.002743612 | 0.04872248 | -1.48070 |
| TBC1D8-AS1 | 1.33224 | 0.31838 | 3.64369 | 0.002744311 | 0.04872248 | -1.42658 |
| HHLA1 | 1.39204 | 2.31567 | 3.64846 | 0.002755896 | 0.048876102 | -1.54321 |
| SNHG7 | -0.47084 | 3.81627 | -3.63906 | 0.002769337 | 0.049062292 | -1.86961 |
| MEG8 | 1.29381 | 3.38873 | 3.64139 | 0.002794212 | 0.049450439 | -1.73107 |
| ADH1B | 1.60043 | 1.94365 | 3.63299 | 0.002802522 | 0.049544903 | -1.45353 |
| TAS2R20 | 1.79782 | -0.48711 | 3.63196 | 0.002808188 | 0.049592477 | -1.50286 |
| LINC01182 | 1.64620 | 0.88312 | 3.62903 | 0.002824388 | 0.04982579 | -1.45221 |
| PTN | -1.63252 | 0.43443 | -3.63092 | 0.002834232 | 0.0499249 | -1.48781 |
| TERF1 | 0.80755 | 5.04673 | 3.63349 | 0.002837705 | 0.0499249 | -2.02436 |
| RPS12P16 | 2.13234 | -1.17528 | 3.62640 | 0.00283899 | 0.0499249 | -1.62356 |
